# Boosting Methane Production at Low Temperatures: The Role of Zero Valent Iron-Driven Syntrophy in Anaerobic Digestion

**DOI:** 10.64898/2026.11.24.768983

**Authors:** Chenxiao Liu, Dominic Frigon

## Abstract

The efficiency of anaerobic digestion (AD) declines at low temperatures due to reduced microbial metabolic rates and weakened syntrophic interactions, prompting the need for strategies that stimulate syntrophic metabolism between acetogens and methanogens. This study investigated whether the enhancement of syntrophic metabolism by zero valent iron (ZVI) under psychrophilic conditions (20 °C) depends on the electrical conductivity of the particles, as opposed to the biofilm-carrier effects shared by any retained particle. A non-conductive silicon dioxide-coated ZVI (ZVI@SiO_2_) was synthesized and used as a biofilm-carrier control that retains the particle surface and retention time but electrically isolates the iron core. Long-term semi-batch reactors revealed that ZVI significantly enhanced methane production, volatile solids destruction, and volatile fatty acid (VFA) conversion, outperforming both non-particle control and ZVI@SiO_2_ control. Batch incubations with methanogenic inhibitors, applied to biofilm-only and bulk-only inocula, localized the enhancement to ZVI particle-associated biofilms. RNA- and DNA-based microbial community analysis showed that ZVI promoted the activity of syntrophic bacteria (e.g., genera *Smithella* and *Syntrophomonas*) and hydrogenotrophic methanogens (e.g., genera *Methanoregula* and *Methanolinea*), supported by elevated *mcrA* gene expression on ZVI biofilms relative to ZVI@SiO_2_. Acetoclastic *Methanothrix* remained the most abundant methanogen genus, while hydrogenotrophic *Methanoregula*, *Methanolinea*, and *Methanospirillum* exhibited selectively elevated *mcrA* mRNA-to-DNA ratios on ZVI biofilms, indicating coexistence and transcriptional activation of both acetoclastic and hydrogenotrophic pathways within the ZVI biofilm niche. The enhancement was lost when the iron core was electrically insulated, and the enriched taxa and transcriptional patterns followed the trends reported for systems in which direct interspecies electron transfer (DIET) has been demonstrated.

## 1. Introduction

There is great interest in low-temperature (psychrophilic) anaerobic digestion (AD) (e.g., 10– 20 °C) to save energy, especially in cold climates, but this comes with significant challenges. Digester heating is a major component of the energy demand of sludge digestion, and the net energy benefit of operating ozone-pretreated WAS digesters at low temperature has been quantified previously^1^; any additive that recovers part of the kinetic penalty of low temperature therefore has a direct energy value. While cold-adapted syntrophic bacteria and methanogenic archaea can grow at these psychrophilic temperatures making feasible methane production^2^, there are significant kinetic limitations to adapt high-rate AD to such conditions. These limitations throughout the entire food web of anaerobic digestion range from disintegration and hydrolysis to electron transfer between acetogenesis and methanogenesis. Ozonation has been shown to effectively alleviate the bottlenecks associated with disintegration and hydrolysis by enhancing solubilization of organic matter at low temperatures^1, 3^. However, the accumulation of volatile fatty acids (VFAs) indicates that the final step of electron transfer—critical for syntrophic metabolism—remains a key limiting factor at low temperatures^3^. This bottleneck has largely been attributed to the slow diffusion rate of mediated interspecies electron transfer (MIET) via acetate, hydrogen, and formate^4^.

A growing body of research suggests that additions of conductive material, including magnetite, granular activated carbon (GAC), biochar, and zero-valent iron (ZVI), can enhance direct interspecies electron transfer (DIET), leading to improved methane production^5^. By opposition to MIET, DIET is the transfer of higher energy electrons to methanogens by electroconductive materials forming bridges with the syntrophic consumers of fatty acids. Evidence for DIET facilitated by conductive materials was first demonstrated in a coculture of *Geobacter metallireducens* and *Methanosarcina barkeri* using (GAC), with ethanol supplied as the sole electron donor^6^. More complex applications of conductive materials, including ones using ZVI, were investigated across various waste types, where enhanced methane production was consistently observed^7^. The improved methanogenesis kinetics were attributed to enhanced DIET between syntrophic bacteria and methanogens^8^. If such a route operates in complex sludge communities, it would be most valuable under low-temperature conditions where diffusion-limited MIET is slowest. However, in digesters the contribution of DIET is generally inferred from community composition and performance proxies rather than measured directly, and the enhancement obtained with conductive additives may also arise from non-conductive effects such as biofilm formation, surface adsorption, and the longer retention of particle-attached biomass. Separating these contributions requires a control that preserves the particle but removes its conductivity, which has rarely been done in ZVI-amended digesters.

The current study focused on revealing the mechanisms leading ZVI to enhance methane production during the low temperature high-rate AD of waste activated sludge (WAS). A first hypothetical mechanism is the anaerobic abiotic oxidation of ZVI’s releasing hydrogen to support hydrogenotrophic methanogenesis^9^. However, stoichiometric calculations and complementary abiotic experiments suggested that these mechanisms contributed less than 1% of the methane production enhancement^10^. A second hypothetical mechanism is the formation of biofilm at the surface of ZVI particles due to their hydrophobicity and porosity^11^. In the system under study here and previously, the digestate solids retention time was 20 days, while the ZVI retention times was 60 days, which could contribute to the enrichment of slow-growing methanogenic archaeal and associate syntrophic bacterial populations. Furthermore, the close association of the population in the dense biofilms may reduce diffusion limitations to enhance MIET and improve electron exchange (i.e., DIET) through structures like e-pili^12^. A final hypothetical mechanism relies on the high electrical conductivity of ZVI to transport electron through relatively long distances between syntrophs and methanogens^13^.

These hypothetical mechanisms were investigated here by first operating a new control reactor system with silicon dioxide-coated ZVI particles (ZVI@SiO_2_). These particles can be manipulated (magnetic separation, retention) in a fashion similar to the native ZVI particles and offer a comparable surface for biofilm development, but their iron core is electrically insulated. Any enhancement shared by ZVI and ZVI@SiO_2_ can thus be attributed to biofilm-carrier effects, whereas the enhancement specific to ZVI isolates the conductivity-dependent fraction. Then microbial enrichment was examined by sequencing of both 16S rRNA and rRNA gene, and of the functional gene *mcrA*, a robust marker of methanogenic potential due to its strong correlation with methane production rates^14^. Understanding microbial community dynamic is essential for elucidating ZVI’s impact on low-temperature AD. While DNA-based techniques reveal the total microbial community, they fall short in identifying the most metabolically active populations. RNA-based profiling overcomes this limitation by revealing the most transcriptionally active taxa; notably, DNA- and RNA-based profiles in full-scale anaerobic digesters have been reported to differ markedly, with 26% of bacterial and 42% of archaeal OTUs showing marked differences in relative abundances^15^.

Beyond community composition profiling, specific inhibitors of methanogenic metabolism were employed to further delineate the role of syntrophic bacterial species and methanogenic archaeal species in response to ZVI addition, and to distinguish the role of hydrogenotrophic and acetoclastic methanogenesis. Fluoroacetate is a specific inhibitor of acetate metabolism. It is metabolised into fluorocitrate in the first step of the TCA cycle, which blocks the next step catalysed by aconitase^16^. Despite the TCA cycle not being involved in acetoclastic methanogenesis, fluoroacetate likely interferes with their metabolism downstream of acetyl-CoA synthesis, and it is widely used to differentiate between acetoclastic and hydrogenotrophic pathways^17, 18^.

In parallel, chloroform, a broad-spectrum methanogenic inhibitor, was used as an overall control to suppress methanogenesis more generally. Chloroform inhibits both acetoclastic and hydrogenotrophic methanogens by targeting corrinoid enzymes and methyl-coenzyme M reductase^19^.

The novelty of this work lies not in reconfirming that ZVI increases methane yield, which is well established^7^, but in (a) separating the conductivity-dependent fraction of the enhancement from the biofilm-carrier effects with an electrically insulated ZVI control, (b) localizing the enhancement to the particle-associated biofilm by inhibitor-dissected incubations of biofilm-only and bulk-only inocula, and (c) resolving the active taxa by paired RNA- and DNA-based 16S rRNA and *mcrA* profiling under psychrophilic conditions. We therefore test two complementary hypotheses: (i) the methane enhancement conferred by ZVI under low-temperature AD is conductivity-dependent, lost when the conductive iron core is electrically isolated by a non-conductive SiO_2_ coating, and (ii) the taxa and transcriptional responses that track this conductivity-dependent enhancement follow the trends reported for systems in which direct interspecies electron transfer (DIET) has been demonstrated, which would support DIET as a hypothesized, though not directly tested, contributor within the syntrophic biofilm developed on the particle surface.

## 2. Material and methods

### 2.1. Synthesis of silicon dioxide coated zero valent iron particle

SiO_2_ coated zero-valent iron particles (ZVI@SiO_2_) were synthesized following Wu et al., Wu, Sun, Lu, Cheng and Gao ^20^ with modifications. Briefly, iron powder was cleaned and coated via a TEOS-based microemulsion; detailed methods are provided in the Supplementary Methods 2.1.1. The quality of silicon dioxide coating was examined by focused ion beam and scanning electron microscope (FIB-SEM) dual beam system (FEI Helios Nanolab 660, Hillsboro, USA); detailed methods are provided in the Supplementary Methods 2.1.2. Apparent DC resistivity of packed ZVI and ZVI@SiO_2_ powder was measured (Supplementary Methods 2.1.3).

### 2.2. WAS sampling and long-term operation of anaerobic digesters

The inoculum for long-term anaerobic reactor study was obtained from full-scale mesophilic anaerobic digesters at the water resource recovery facility of the Régie de l’Assainissement des Eaux du Bassin La Prairie in St-Catherine (Québec), Canada. The WAS used to feed these digesters was collected bi-weekly at the same facility. Ozonated at 0.005 g-O_3_/g-VS was applied to the WAS to increase the hydrolysis as descried according to LinLin ^3^.

To conduct the experiment, semi-batch reactors were designed using 500-mL serum bottles, each with a working volume of 400 mL and a headspace of 100 mL. The reactors were fed with ozonated WAS at a concentration of 10 g-TS/L and were maintained at a solids retention time (SRT) of 20 days. Six experimental setups were tested, incorporating two different temperature conditions (20°C and 35°C) and three particle dosing scenarios: a control with no particles, 5 g/L of ZVI@SiO_2_ (biofilm control), and 5 g/L of ZVI. A total of 20 parallel reactors were operated to test the experimental conditions. For the particle dosing conditions, four replicate reactors were operated, while for the no-particle control conditions, only duplicate reactors were operated. Particles were separated from the WAS using a neodymium magnet (McMaster-Carr, USA) and returned to the digesters; the wastage of ZVI particles was adjusted such that their retention time was set at 60 days. The reactors were operated for a total of 144 days to reach steady state. Biogas production and pH were monitored every two days. Weekly analyses of the digestate characteristics included total suspended solids (TSS), volatile suspended solids (VSS), soluble chemical oxygen demand (COD), volatile fatty acids (VFAs), and biogas composition (Section 2.4.). All biomass and particle with biofilm were collected at the end for metagenomic analysis.

### 2.3. Batch incubation with microbial pathway inhibitors

Batch experiments were conducted in 60-mL serum bottles with a 40 mL working volume. Inocula were sourced from the long-term operation of semi-continuous 500 mL anaerobic reactors operated at steady-state at 20 °C (Section 2.2.). Three bulk sludge inocula (20 mL each) were taken from reactors treated with ZVI@SiO_2_, ZVI, and a control. For particle inocula, ZVI@SiO_2_ and ZVI particles were magnetically separated from their respective reactors and added at 0.5 g per bottle. Corresponding supernatants were centrifuged at 5000 rpm for 30 minutes, with 20 mL added to bulk sludge-inoculated bottles and 40 mL to particle-inoculated ones. Each inoculum type was tested in duplicates under three conditions: with fluoroacetate (1 mM), chloroform (100 µM), and a no-inhibitor control, resulting in 30 reactors. Ethanol was added as the sole substrate at an initial concentration of 35 mM. The experimental conditions for the inoculum and inhibition are summarized in Table S1. All bottles were purged with high-purity nitrogen for 10 min before inhibitor addition to ensure anaerobic conditions. Reactors were incubated at 20 °C and monitored every 2 days for the first 8 days, then every 4 days until day 28. Bulk sludge-inoculated reactors were operated for 38 days, and particle-inoculated reactors for 62 days. Biogas production, gas composition, pH, and VFAs were measured throughout. An electron-equivalent and carbon balance was computed for each reactor from ethanol input, initial biomass, VFA accumulation, CH_4_, CO_2_ and H_2_ production, and endpoint biomass (Supplementary Methods 2.5.1). All biomass and particle with biofilm were collected at the end for metagenomic analysis.

### 2.4. Physical and chemical analytic method

Biogas production was monitored using a pressure gauge (McMaster-Carr, USA) and converted into volume. The pH of the digestate was measured using an Orion pH meter (ThermoFisher Scientific, USA). Weekly analyses of TSS (Method 2540 D), VSS (Method 2540 E), and COD (Method 5220 D) were conducted according to standard methods APHA et al., ^21^. Ethanol and VFAs were quantified using a 7820 A gas chromatograph (Agilent Technologies, USA) with a flame-ionization detector (FID). The headspace gas composition was also determined using the 7820 A gas chromatograph with a thermal conductivity detector (TCD) for methane and carbon dioxide measurements^3^. Steady-state was defined as stable reactor operation for at least one SRT.

### 2.5. 16S rRNA and mcrA amplicon sequencing

For long-term reactors dosed with ZVI@SiO_2_ or ZVI particles, the digestates were magnetically separated into bulk sludge and particles with biofilm. The particles with biofilm were washed three times with deionized water, and the biofilm was then collected by physically exfoliating the particles using a Precellys®24 tissue homogenizer (Bertin Technologies, France). The biomass (bulk sludge-inoculated reactors) and particle with biofilm (particle inoculated reactors) from batch incubation with inhibitors were also collected at the final stage for DNA and RNA extraction. RNA was extracted from both the bulk sludge and biofilm samples using the RNeasy PowerSoil Total RNA Kit (Qiagen, USA). During the RNA extraction process, DNA was also isolated using the RNeasy PowerSoil DNA Elution Kit (Qiagen, USA). DNase I (Invitrogen, USA) was used to remove DNA contamination from RNA samples. The 16S rRNA (V4) and *mcrA* genes were amplified, purified, and quantified from DNA and RNA templates using standard PCR and RT-PCR workflows described in the Supplementary Methods 2.5. The purified libraries were pooled and sequenced on the Illumina NextSeq platform using paired-end 300 bp (PE300) reads at the Génome Québec Innovation Centre (Montréal, Québec, Canada). All raw sequencing data have been deposited in the NCBI SRA under BioProject PRJNA1401889.

### 2.6. Digital PCR

Absolute quantification of *mcrA* and 16S rRNA (housekeeping gene) was performed via dPCR (QIAcuity One, Qiagen, USA) for all reactors DNA and RNA samples. Optimal dilution ratios were determined by qPCR, and samples were digested with EcoRI (0.25 U µL^−1^) prior to nanoplate loading. Detailed dPCR protocols, including qPCR for dilution ratios, imaging parameters and cycling conditions, are provided in Supplementary Method 2.6.

### 2.7. Bioinformatic analysis

Bioinformatics analysis was conducted using R with denoising performed by DADA2 package^22^. Taxonomic classification amplicon sequence variants (ASVs) was assigned using the MiDAS 4.8.1 database for 16S rRNA (and gene)^23^, and using NCBI BLAST for *mcrA* (mRNA and gene). The microbial community structure was further analyzed using the Vegan^24^ and Complex heatmap^25^ package in R. PICRUST 2^26^ was used to predict the metabolic pathway.

For statistical analysis, one-way ANOVA was performed using the default function in R to assess the significance of difference across the different operational conditions.

## 3. Results

### 3.1. Silicon dioxide coating

The quality of silicon dioxide coating was imaged using a focused ion beam and scanning electron microscope (FIB-SEM) with EDS elemental mapping. Cross-sectional imaging revealed iron cores surrounded by a coherent SiO_2_ layer (Fig. 1 and S1). The elemental mapping of silicon and oxygen using both cross-sectional and surface images consistently overlapped and displayed the relative abundance ratios expected for SiO_2_, confirming the presence of SiO_2_ as the outer layer (Fig. S1 and S2). The coating thickness averaged 242.8 ± 82.3 nm (median 239.6 nm), consistent with previous reports for similar core–shell particles^20, 27^. The apparent DC resistivity of the packed ZVI@SiO_2_ was 38-fold higher than that of ZVI (Table S4). All our characterization of the resulting particles confirmed that the SiO_2_ coating provided an amorphous structure that prevented oxidation of the ZVI surface and blocked the direct electron transfer through the conductive ZVI core.

**Fig. 1.**
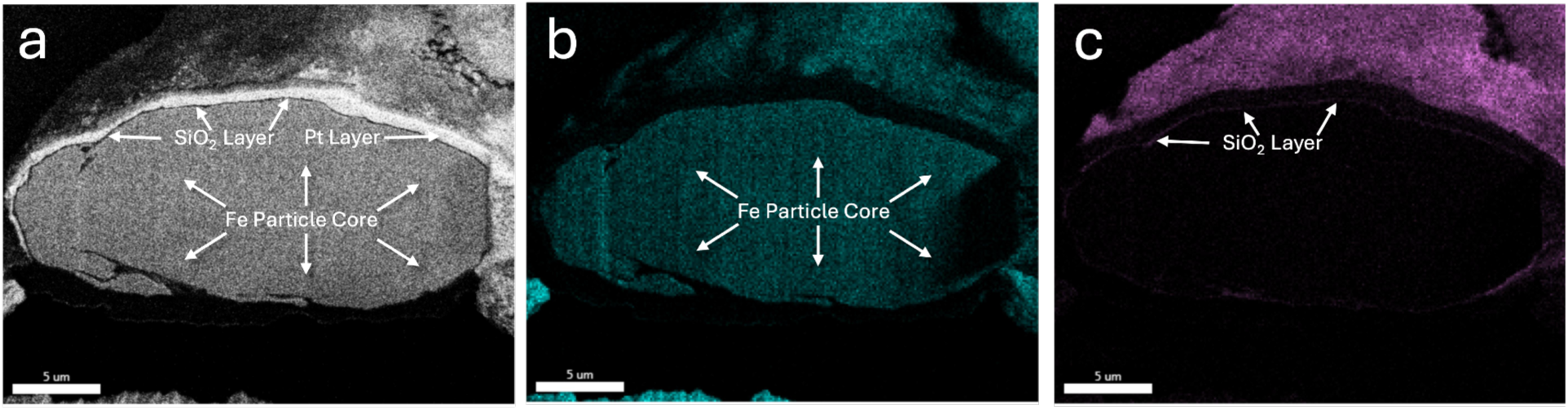
Cross-sectional FIB-SEM imaging and EDS elemental mapping of silicon oxide coated ZVI (ZVI@SiO₂) particles. a) Complete image of the particle cross-section, the outer layer (white) is a 2 μm thick platinum layer to protect the particle structure during the milling preparatory process, the SiO_2_ layer is observable in black between the iron core and the platinum layer; b) EDS elemental mapping of Fe (teal colour); c) EDS elemental mapping of Si (magenta colour).

### 3.2. Long-term performances of digesters

During the long-term operation of digesters, reactors amended ZVI particles produced significantly higher methane yields (p≤0.05) than both the no-particle control and non-conductive ZVI@SiO_2_ particle control. Methane production increased by 21% to 28% at 20 °C and 5% to 17% at 35 °C depending on the criterion used for comparison absolute methane productions yield (Table S5), specific methane production normalized to volatile suspended solids fed (SMP_VSS,in_; Table S5), or specific methane production normalized to volatile suspended solids destroyed (SMP_VSS,des_; Fig. 2a). Conversely, reactors dosed with ZVI@SiO_2_ showed methane production levels similar (p>0.05) to the control reactors across all three criteria at both temperatures. The average VSS destruction was generally higher at 35 °C than at 20 °C, likely due to increased enzymatic activity at the higher temperature, promoting greater solids consumption and electron transfer into methane. Otherwise, VSS destruction rates aligned with methane production trends, reaching the highest levels in ZVI reactors at both temperatures (Fig. 2b). Conversely, ZVI reactors showed the lowest accumulation of VFAs at both temperatures (Fig. 2d), which likely contributed to the significant increase in methane production. Additionally, the VFAs profile in the ZVI reactors was dominated by acetic acid, in contrast to the accumulation of longer-chain VFAs observed in the other reactors. Finally, methane proportions in the gas phase were not significantly different (p>0.05) between reactors receiving different particle dosage, but were higher at 20 °C than at 35 °C (Fig. 2c), probably due to CO_2_ solubility being reduced with increasing temperatures.

**Fig. 2.**
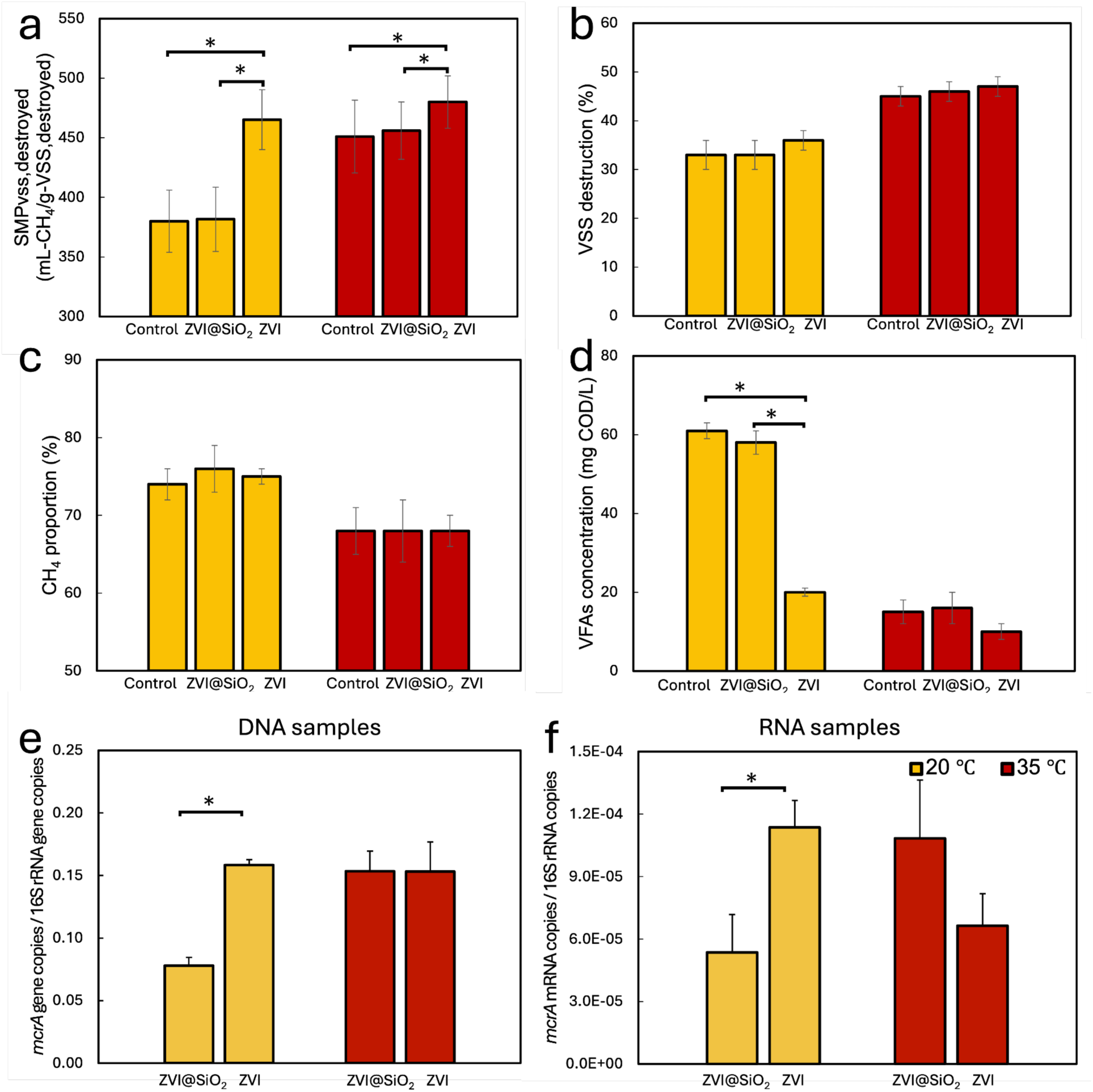
Performance of anaerobic digestors with ZVI@SiO2 and ZVI addition (5 g/mL) at 20 °C (yellow bars) and 35 °C (red bars). a) specific methane production normalized to VSS destroyed (SMPvss,des); b) VSS destruction; c) CH_4_ proportion; d) total VFA concentrations. And relative quantification of *mcrA* in particle surface biofilms normalized by their 16S rRNA abundances at 20 °C and 35 °C. e) *mcrA* gene relative abundance in DNA extracts; f) *mcrA* transcripts relative abundances in RNA extracts. Asterisks indicate significantly different statistical comparison pairs: p ≤ 0.05 (*). Error bars represent standard deviations of biological replicates (n = 2 for non-particle controls, n = 4 for particle treatments).

The relative abundance of *mcrA* gene and *mcrA* mRNA (normalized to16S rRNA gene copies and 16S rRNA copies, respectively) for samples from particles’ surfaces (Fig. 2e and f) and bulk sludges (Fig. S3a, b) were quantified through digital PCR approach. At both temperatures, *mcrA* gene relative abundance was consistently higher at the surface of particles compared to the bulk, indicating a clear enrichment of methanogens on particle surfaces. Similarly, the relative activity of methanogens, represented by the *mcrA* mRNA to 16S rRNA ratio, was generally higher on particle surfaces than in the bulk, suggesting elevated methanogenic activity. At 20 °C, *mcrA* gene relative abundances were on average doubled on ZVI particles (0.158 *mcrA*/16S rRNA gene copies) from the ones on ZVI@SiO_2_ particles (0.078; Fig. 2e). Conversely, the *mcrA* gene relative abundances on the surfaces of both particle types were similar at 35 °C. A similar pattern was observed for *mcrA* mRNA, with higher expression on ZVI surfaces at 20 °C (Fig. 2f). However, the relative expression of *mcrA* appeared to have dropped at 35 °C on ZVI surfaces compared to ZVI@SiO_2_ surfaces, but the difference was not significant (p ≥ 0.05). In the bulk sludges, *mcrA* gene relative abundances were highest at 35 °C in control condition, but the difference with the other reactors was only significant with the ones receiving ZVI at 35 °C (Fig. S3a).

### 3.3. Microbial diversity of long-term reactors under different treatments

#### 3.3.1. Alpha diversity of long-term anaerobic digesters

ZVI particles harboured communities with higher first-order Hill’s numbers (derived from Shannon diversity) than the communities in the bulk solids of the same reactors for 16S rRNA (and gene) and *mcrA* amplicon sequencing analyses of both RNA and DNA (Fig. S4), suggesting greater species diversity^28^. The only exception was observed for *mcrA* diversity at 35 °C (Fig. S4c). In contrast, ZVI@SiO_2_ particle amended reactors showed opposite trends in first-order Hill’s numbers diversity when comparing communities on particle surfaces and the bulk of the same reactor for all but two of the analyses.

#### 3.3.2. Differences in active bacterial population structures in long-term anaerobic digesters

The structure of the active bacterial community (RNA-based) was primarily driven by location (bulk vs. particle surface), followed by temperature (20 °C vs. 35 °C), and then treatment (digesters receiving ZVI particles vs. ZVI@SiO_2_ particles vs. control; Fig. 3a). In contrast, the archaeal population assembly was first driven by temperature, with secondary effects of location and treatment (Fig. 3b).

**Fig. 3.**
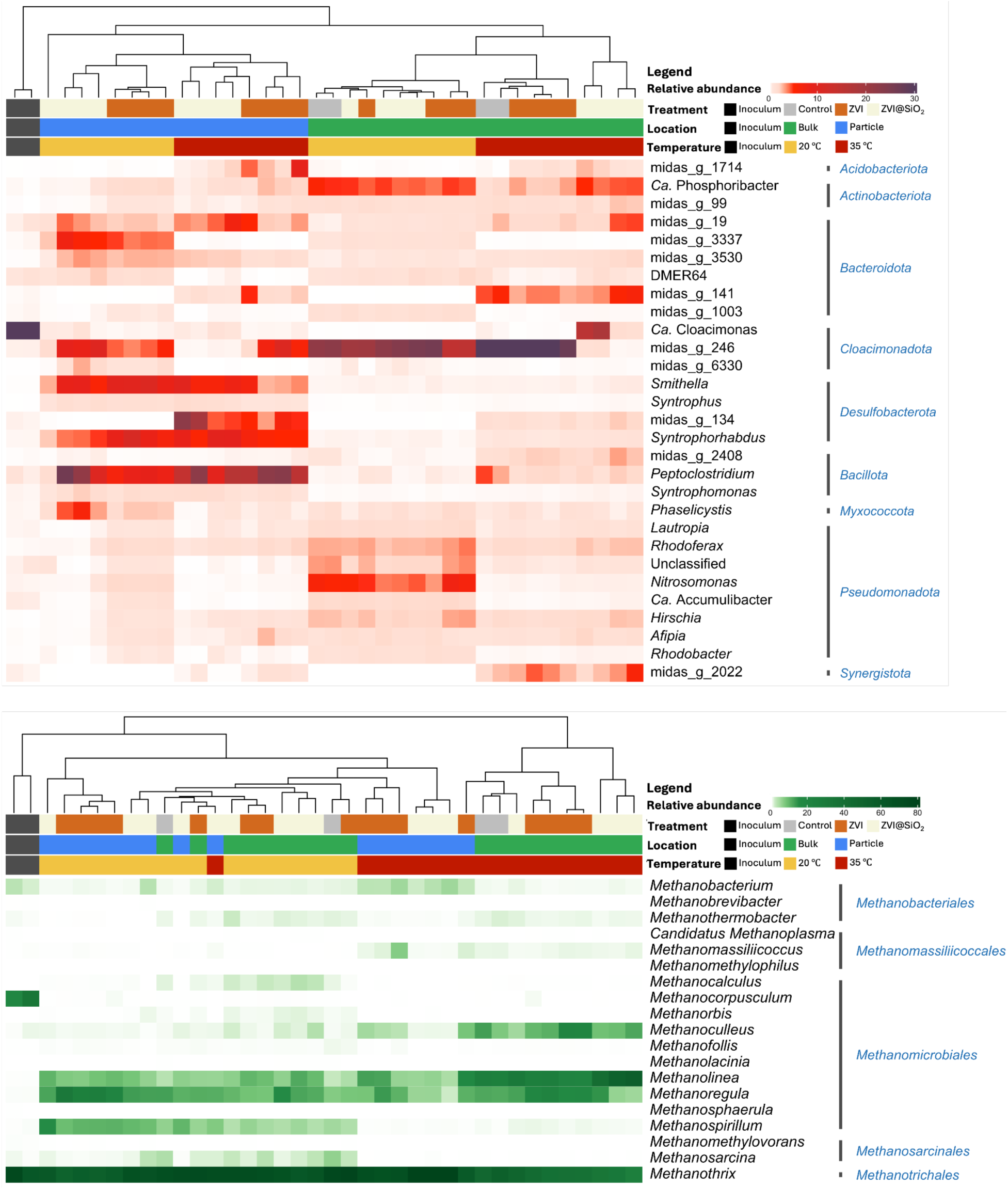
Beta-diversity dendrogram and heatmap of the relative abundances of genera in anaerobic digestors receiving ZVI or ZVI@SiO_2_ at 20 or 35 °C determined by 16S rRNA (a) and *mcrA* mRNA (b) RT-amplicon sequencing. The dendrogram is based on Bray-Curtis dissimilarity calculated from amplicon sequence variants (ASVs) and clustered by UPGMA method. The heatmap includes the top 15 most abundant genera of each sample for both 16S rRNA and *mcrA* mRNA. For 16S rRNA, taxonomic annotations are shown from left to right as genus and phylum (in blue). For *mcrA* mRNA, annotations are shown as genus and order (in blue). All methanogenic genera shown for the *mcrA* mRNA panel belong to the phylum *Euryarchaeota*. Relative abundances are represented by color intensity, with red for the 16S rRNA and green for the *mcrA* mRNA. Reactor conditions are color-coded based on treatment (Control, Inoculum, ZVI, ZVI@SiO_2_), location (Bulk, Particle), and temperature (20 °C, 35 °C).

Location was the primary driver shaping the active bacterial community (as represented by RNA profiles), likely influenced by the longer retention of particles (60 d) relative to the digestate (20 d) and by the development of specific micro-niches at the particle surface, which favoured microbial colonization and growth on both particle types irrespective of their conductivity. Particle surfaces were dominated by *Peptoclostridium* (phylum *Bacillota*; Fig. 3a), and at 20 °C, showed enrichment of midas_g_3337, midas_g_3530 (both in phylum *Bacteroidota*), *Smithella*, and *Syntrophorhabdus* (both in phylum *Desulfobacterota*). The last two genera showed enrichments in both RNA and DNA extracts (Fig. S5a). In contrast, the active microbial community RNA extracts from the bulk sludge were different from the particles surfaces by their elevated abundances of Candidatus *Phosphoribacter* (phylum *Acidobacteriota*), and midas_g_246 (phylum *Cloacimonadota*), with *Nitrosomonas* (phylum *Pseudomonadota*) being specifically enriched at 20 °C. At 35 °C, the bulk-associated communities shifted, showing enrichment of midas_g_141 (phylum *Bacteroidota*) and midas_g_2022 (phylum *Synergistota*).

The DNA extracts revealed both similar and divergent trends to the one found with the RNA extracts. At 20 °C, the DNA extracts showed higher abundances in bulk biomass of Christensenellaceae_R-7_group, *Sedimentibacter* (both in phylum *Bacillota*), DMER64, midas_g_12 (both in phylum *Bacteroidota*), midas_g_249 (phylum *Synergistota*), and midas_g_467 (phylum *Chloroflexota*) (Fig. S5a), while particle-associated communities were characterized by *Syntrophomonas*, *Cryptanaerobacter* (both in phylum *Bacillota*), midas_g_762 (phylum *Synergistota*), and midas_g_3337 (phylum *Bacteroidota*).

#### 3.3.3. Differences in active archaeal methanogenic population structures in long-term anaerobic digesters

At 20 °C, *Methanothrix*, *Methanospirillum*, and *Methanoregula* were generally more abundant comparing to 35 °C, with *Methanospirillum* and *Methanoregula* specifically enriched in the particle-associated communities (Fig. 3b). Among these, *Methanoregula* showed higher abundances on the ZVI particle surfaces. In contrast, *Methanothrix* was more prevalent in the bulk phase at 20 °C, whereas at 35 °C, *Methanothrix* became more enriched on particle surfaces. Additionally, *Methanolinea* and *Methanoculleus* were more abundant overall at 35 °C, with both *Methanoregula* and *Methanoculleus* particularly enriched in the bulk phase of ZVI reactors at this higher temperature.

### 3.4. Probing community food-web with methanogenesis inhibitors

#### 3.4.1. Performance of batch reactors

To better understand the mechanisms underlying enhanced methanogenic activity associated with ZVI particles, a series of batch incubations were conducted with the addition of specific inhibitors to dissect the microbial pathways involved. Building on the hypothesis that ZVI addition could influence syntrophic bacteria and methanogens, and to distinguish between hydrogenotrophic and acetoclastic methanogenesis, inhibition assays were conducted. These incubations also tested whether the conductivity-dependent advantage of the ZVI biofilm persisted when acetate metabolism, or methanogenesis as a whole, was inhibited. Ethanol was provided as the sole substrate, and bulk sludge and particle-associated biofilms were incubated separately under control and inhibitor treatments to evaluate their respective contributions to methane production.

The results from the batch incubations showed that ZVI particle-inoculated control reactors (without inhibitors) produced significantly higher cumulative methane than those inoculated with ZVI@SiO_2_ particles (Fig. 4a, b), which is reminiscent of the long-term reactor performances. This trend corresponded with more efficient VFA degradation in ZVI particle-inoculated control reactors, whereas acetic acid and propionate acid accumulated in ZVI@SiO_2_ particle control reactors (Fig. 4c, d). In contrast, methane production did not significantly differ among bulk sludge-inoculated reactors regardless of the inoculum source (20 °C long-term reactors treated with ZVI, ZVI@SiO_2_, or control; Fig. S6a, b, c). Under fluoroacetate inhibition, cumulative methane production in both particle-inoculated reactors (ZVI and ZVI@SiO_2_) plateaued at similar levels, and their VFA profiles paralleled each other for the accumulation of acetic acid and propionate acid (Fig. 4c, d). Finally, chloroform addition essentially eliminated methane production (Fig. 4a, b), but led to the production of longer chain VFAs (Fig. 4c, d), which is likely the result of chain elongation acting as electron acceptor^29^. These chain elongation processes, including ethanol-to-propionate conversion via the acrylate pathway, are known to occur across diverse anaerobic taxa^30^. Although, it was notable that this last activity was much delayed for the biomass on the ZVI particles.

**Fig. 4.**
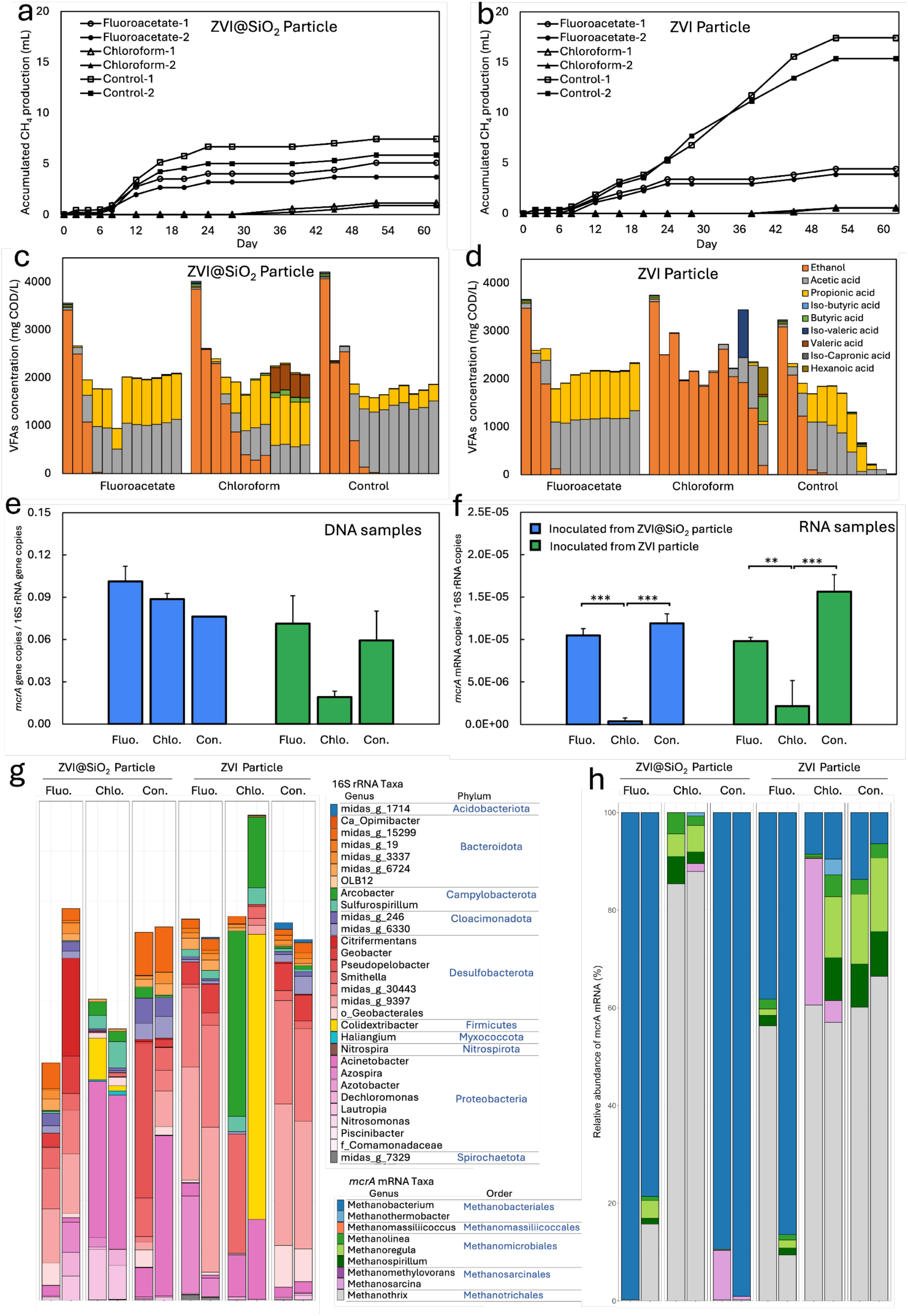
Temporal accumulated CH_4_ produced (a,b) and VFAs concentration (c,d) profiles under various inhibitor additions in batch reactors at 20 °C inoculated with particles from long-term digesters at the same temperature: (a,c) ZVI@SiO_2_ particles, (b,d) ZVI particles. At the end of the incubation period, relative quantification of *mcrA* gene (e) and mRNA (f) and community profiles determined with RNA extracts for bacteria (16S rRNA, g) and methanogenic archaea (*mcrA*, h). For 16S rRNA (g), only the overall most abundant 30 bacterial genera are presented and the methanogenic genera were removed. Abbreviations: Con. for control reactors without addition of inhibitor, Fluo. for fluoroacetate, Chlo. for chloroform. Taxonomic annotations are shown at the genus and phylum (in blue) levels for 16S rRNA, and at genus and phylum (in blue) levels for *mcrA* mRNA. Significant pair-wise differences: *, p ≤ 0.05; **, p ≤ 0.01; ***, p ≤ 0.001. Error bars represent standard deviations of biological replicates (n = 2).

An electron-equivalent balance across the reactors partitioned the input pool (ethanol plus initial biomass) among CH_4_, VFAs, H_2_, and end biomass electrons (Table S6). Under no-inhibitor conditions, the conductive ZVI particle biofilm converted 26.2 % of the input electrons to CH_4_ with only 6.7 % residual as VFAs, while the non-conductive ZVI@SiO_2_ particle biofilm converted only 11.4 % to CH_4_ and retained 43.1 % as VFAs — a 2.3-fold conductivity-dependent CH_4_ advantage. Under fluoroacetate this advantage collapsed (6.6 % vs. 7.5 % CH_4_ on ZVI vs. ZVI@SiO_2_ particle biofilms).

#### 3.4.2. mcrA gene relative abundance and methanogenesis activity

In inhibitor-treated reactors, fluoroacetate-inhibited reactors showed *mcrA* gene abundances similar to their respective controls across all inoculum types (Fig. 4e), including the bulk sludge where differences appeared more pronounced but were not statistically significant (Fig. S7a). At the RNA level, *mcrA* mRNA expression was likewise comparable between fluoroacetate and control treatments (Fig. 4f, Fig. S7b), in contrast to the significant drop observed under chloroform treatment.

#### 3.4.3. Community diversity and structure of batch reactors

Relative abundance of 16S rRNA (Fig. 4g) revealed that *Geobacter* (phylum *Desulfobacterota*) reached 4.98% and 4.59% (mean duplicates) in ZVI particle-inoculated reactors under control and fluoroacetate conditions, respectively, representing up to a 119-fold increase compared to the initial inoculum. While *Smithella* (phylum *Desulfobacterota*) was highly abundant (6.98%) in the ZVI particle-inoculated control reactor, it showed no enrichment under other conditions. For *mcrA* gene (Fig. S9b) and *mcrA* mRNA relative abundances (Fig. S10b), the number of detected methanogenic genera decreased from 15 (DNA extracts) to 9 (RNA extracts), supporting the observed reduction in Hill’s number (Supplementary Results 3.4.3; Fig. S8) and indicating fewer transcriptionally active methanogens. Under fluoroacetate inhibition, methanogenic activity was concentrated in a smaller group, with *Methanobacterium* comprising up to 89.2% in particle-inoculated reactors and 60.4% in bulk sludge-inoculated reactors paralleled by a decline of *Methanothrix* that dominated the inoculum. This pronounced shift from *Methanothrix* to *Methanobacterium* dominance was observed specifically in the ZVI@SiO_2_ particle-inoculated control reactors.

## 4. Discussion

### 4.1. Long-term reactor performance and methanogenic activity

In the long-term experiment, we examined whether enhanced methane production with ZVI addition was driven by particle conductivity or other particle-associated effects such as extended SRT. The addition of ZVI to the reactors clearly increased the methane production above both the no-addition control and the ZVI@SiO_2_ addition reactors (Fig. 2a). This was due to improved COD and VFA removal efficiencies, but not an increase in VSS destruction. Because the non-conductive ZVI@SiO_2_ particles did not reproduce this enhancement despite comparable biofilm formation, a property of the native ZVI surface that is removed by the SiO_2_ shell, most plausibly its electrical conductivity, is the most parsimonious explanation for the methane gap. This conductivity-dependent contribution follows the same trend as in conductive-material studies where DIET between syntrophs and methanogens has been invoked. We therefore regard DIET as a hypothesized contributor on the particle surface; our data do not demonstrate electron transfer through the particles, and the non-conductive effects of the biofilm niche (extended SRT, reduced diffusion distance) that are shared by both particle types operate in parallel.

The observed increase in efficiency of COD conversion into methane by the addition of conductive material is consistent with previous findings about anaerobic digestion. For example, upflow anaerobic sludge blanket reactors operated at 18 °C and supplemented with granular activated carbon (GAC) achieved a 21% increase in methane production and a 20% reduction in effluent soluble COD compared to reactors adding non-conductive sponges^31^. Similarly, anaerobic digesters treating swine wastewater produced more methane when amended with granular graphite or biochar, with increases of 23 ± 11% and 12 ± 6.1%, respectively, compared with non-conductive glass particles or no-particle controls^32^.

The increase in both the relative abundance and expression of the *mcrA* gene in long-term reactors corresponded well with the observed methane production rates at 20 °C. With the application of digital PCR with methanogen (*met* and *mcrA*) gene, similar correlation between gene absolute quantities and their methane production has been be demonstrated dynamically^33^. In current study, higher *mcrA* gene relative abundance and expression were consistently observed on particle surfaces compared to the bulk, with further enhancement on conductive ZVI particles relative to non-conductive ZVI@SiO_2_ at 20 °C. At 35 °C, differences between treatments were less pronounced and not statistically significant, likely because mesophilic conditions already support near-maximal methanogenic activity^34^. Under these conditions, electron transfer may no longer be the rate-limiting factor, thus the conductivity advantage of ZVI becomes less significant. These observations suggest that particles, particularly conductive ones like ZVI, may support higher methanogenic activity, especially under low-temperature conditions.

### 4.2. Microbial taxa adapted to low-temperature anaerobic digestion

Comparison of the community profiles from both DNA and RNA results revealed several taxa that were enriched under psychrophilic conditions (20 °C) in both the ZVI particle biofilms and the bulk sludge. In particular, midas_g_3337, and midas_g_3530 were enriched on ZVI particle surfaces, while DMER64, *Sedimentibacter*, and *Nitrosomonas* were more abundant in the bulk sludge. These taxa likely represent microbial populations adapted to low-temperature anaerobic digestion. *Nitrosomonas* was unexpectedly enriched in the bulk at 20 °C, likely due to inoculum carryover, micro-oxic niches, or residual rRNA signals from inactive cells^35^. Their absence on ZVI surfaces suggests they are bystanders rather than contributors to the particle-associated methanogenesis.

Functionally, these organisms can be broadly grouped into: 1) Fermenters, which degrade complex organics and supply intermediates, and 2) Syntrophs, which oxidize VFAs in partnership with methanogens. For fermenters, midas_g_3337 and midas_g_3530 (both in phylum *Bacteroidota*) contributed to organic matter degradation and was positively associated with *Methanothrix* in iron-amended systems, enhancing methanogenesis^36^. For syntrophs, such as DMER64 (phylum *bacteroidota*) has been reported to degrade propionate under psychrophilic conditions through syntrophic metabolism, with expressed cold adaptation genes such as *cspA*^37^. Similarly, *Sedimentibacter* and DMER64 may contribute to acetate and hydrogen production in anaerobic systems, supporting downstream hydrogenotrophic and acetoclastic methanogenesis^37^.

### 4.3. Syntrophic interactions and functional adaptations in ZVI-amended anaerobic digestion

To better understand the key microbial interactions stimulated by ZVI, we combined results from two complementary multivariate analyses of communities developed in the two experiments: canonical discriminant analysis (CDA) of the long-term reactors and redundancy analysis (RDA) of the inhibited batch incubations. By examining the overlap between these independent datasets, we can identify microbial populations consistently associated with ZVI surfaces and methanogenic activity, revealing potential core members of the syntrophic network under psychrophilic conditions.

In the CDA of the RNA extracts, 16S rRNA RT-amplicons identified members of 5 genera to be key taxa differentiating the ZVI surface communities from the others at 20 °C (Fig. 5a) with *Smithella*, *Syntrophorhabdu*s, and *Syntrophomonas* being the most abundant ones (Fig. 3a). The same analysis of the *mcrA* mRNA profiles showed that most methanogens showed to be enriched on at 20 °C with the exception of *Methanobacterium* (Fig. 5c). These same bacterial taxa showed higher RNA-to-DNA ratios on the ZVI surface at 20 °C than in the other conditions (Supplementary Discussion 4.3; Fig. S12). Additionally, *Methanoregula* and *Methanolinea* exhibited distinctly high RNA-to-DNA ratio on the ZVI surface biofilm (Fig. S12).

**Fig. 5.**
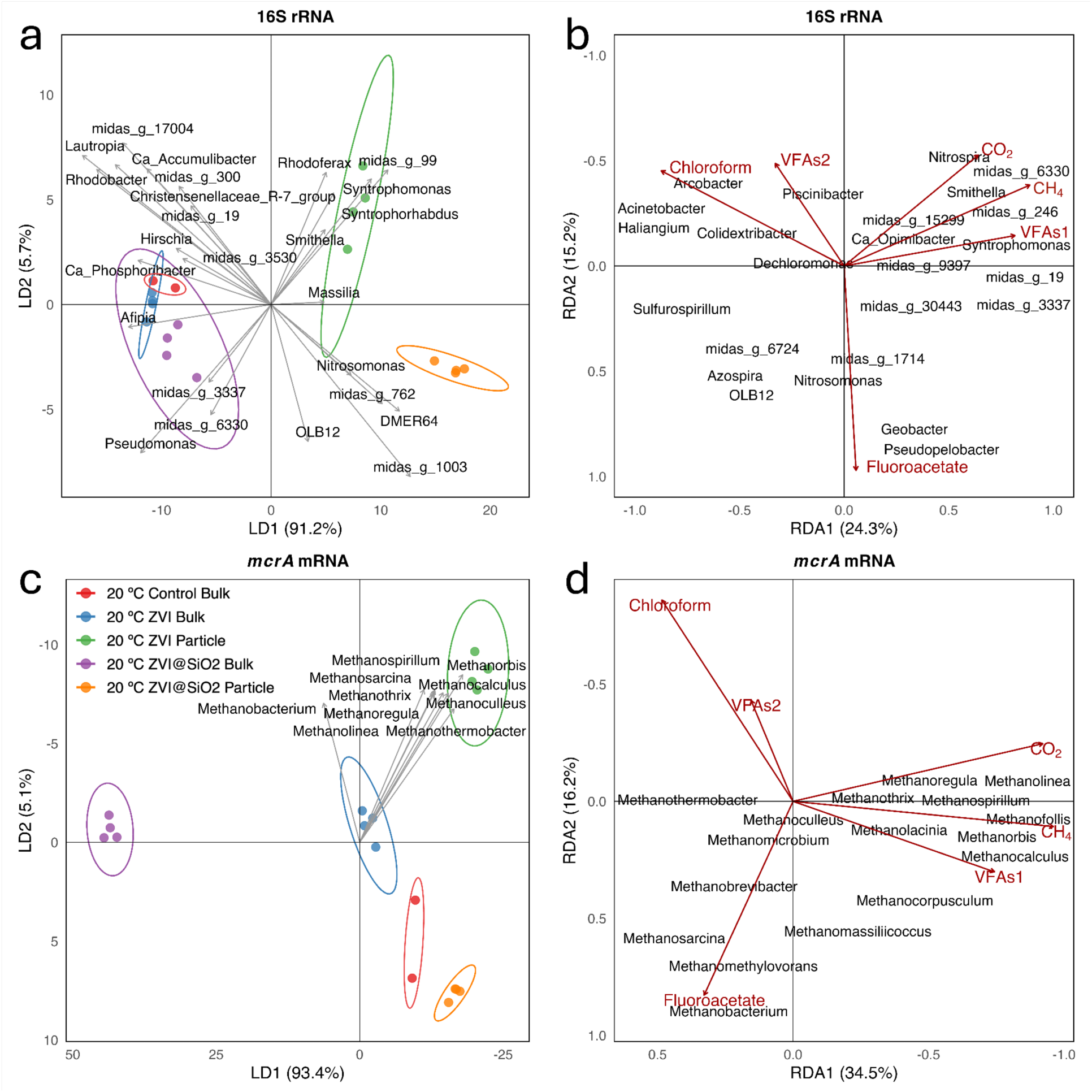
Key microbial taxa at 20 °C determined by RNA extracts and differentiated by long-term reactor conditions (Canonical Discriminant Analysis [CDA] of 16S rRNA bacterial population [a] and *mcr*A mRNA for methanogens [c]) or short-term batch reactors performances with methanogenesis inhibitor additions (Redundancy Analysis [RDA] of 16S rRNA [b] and *mcr*A mRNA [d]). For the RDA, the top 30 genera identified in digesters were included in the analyses. The red arrows represent different performance data and inhibitor additions binary coded as 0/1; note that the arrow no addition control reactors would point along the CO_2_ arrows. Temporal profiles of concentrations in VFAs were reduced in dimensions into two factors together explaining 26% of variability by principal component analysis using Euclidean dissimilarities.

The RDA of the 16S rRNA at the end of the incubations revealed that the natural ZVI-surface community was dominated by members of 7 bacterial genera appeared to be positively correlated with higher methane and CO_2_ production and the principal coordinate 1 of the VFA profiles (VFAs1; Fig. 5b), which captured mainly variations in the acetic acid concentrations. Of these, the genera *Smithella* and *Syntrophomona*s were also enriched on the ZVI surfaces in the long-term anaerobic digesters (Fig. 5a), indicating their importance for the ZVI-associated communities. The 16S rRNA of these populations was less prevalent when the communities were inhibited by either fluoroacetate (acetate metabolism) or chloroform (overall methanogenesis), suggesting that they played a role in the efficient transformation of acetate into methane. Conversely, fluoroacetate addition explained the increase in relative abundances of member of the related genera *Geobacter* and *Ca.* Pseudopelobacter (both in the phylum *Desulfobacterota*). These organisms present in the natural communities on the surfaces of ZVI (Fig. 4c and d) may have consumed the longer chain VFAs that were observed to accumulate with chloroform addition but not acetate nor propionate^38^.

Of the 9 methanogenic genera identified to differentiate the ZVI-associated communities by the CDA of *mcr*A mRNA RT-amplicons, the RDA of the communities described by the same amplicons found correlation of 6 of them with higher methane production: *Methanothix*, *Methanolinea*, *Methanoregula*, *Methanospirillum*, *Methanocalculus*, and *Methanorbis*. Despite this correlation, only the first 4 appeared abundant enough on the ZVI surfaces to warrant further discussion. Two other genera of the original list (*Methanoculleus* and *Methanothemorbacter*) were associated with lower methane production in the batch experiments, and they will not be further discussed. Finally, *Methanobacterium* and *Methanosarcina* appeared associated with fluoroacetate-inhibited samples (Fig. 5b and d). Since fluoroacetate inhibited acetate metabolism, their enrichment likely reflected a shift in the methanogenic community when the natural ZVI-surface assemblage was disrupted, favoring taxa that can utilize alternative substrates under inhibition^39^. These results will be further discussed to highlight the possible syntrophic interactions occurring at the ZVI surface.

Under ZVI supplementation, several syntrophic taxa were consistently enriched, supporting efficient volatile fatty acid (VFA) degradation and methanogenesis. Notably, *Smithella* and *Syntrophorhabdus* were frequently co-enriched and demonstrated strong associations with hydrogenotrophic methanogens such as *Methanoregula*, *Methanolinea*, and *Methanobacterium*, particularly under low-temperature conditions^40^. *Smithella* is recognized for its ability to degrade propionate through a dismutation pathway, producing acetate and butyrate as intermediates^41^. This pathway not only relieves propionate inhibition but also facilitates the growth of syntrophic butyrate degraders. The dominance of *Smithella* in ZVI and Fe_3_O_4_ supplemented systems has been linked to the upregulation of butyrate activation enzymes (e.g., CoA transferase), as well as enhanced DIET-based interactions with methanogens^42^. *Syntrophorhabdus*, on the other hand, was found in the literature to degrade aromatic compounds in cooperation with hydrogenotrophic methanogens through hydrogen transfer. The presence of conductive materials like ZVI and GAC has been shown to strengthen its syntrophic associations, which has been interpreted as enhanced interspecies electron transfer^43^. The co-enrichment of both taxa on ZVI surfaces at 20 °C underscores their functional relevance in maintaining syntrophic metabolism and resilience under psychrophilic conditions. Supporting these syntrophic networks, *Syntrophomonas* contribute to the breakdown of propionate and butyrate into acetate and hydrogen, supplying essential substrates for methanogenesis and reinforcing microbial stability^44^. *Syntrophomonas* clustered phylogenetically near known syntrophic acetate-oxidizing bacteria (SAOB), suggesting potential evolutionary relatedness or functional convergence^45^.

Among methanogens, *Methanothrix* consistently dominated across treatments, particularly in ZVI-amended systems. As a classical acetoclastic methanogen, it benefits from the increased availability of acetate and has been reported to form DIET-capable partnerships with syntrophs such as *Smithella*^42^. Enrichment of *Methanothrix* has been observed in biofilms associated with various ZVI forms (e.g., nano-, strained-, and carbon-supported ZVI), suggesting its adaptability and central role in methane production under both mesophilic and psychrophilic conditions^46^.

Conversely, *Geobacter* species showed varied responses depending on the reactor conditions. Known for their metabolic flexibility and central role in DIET, *Geobacter* spp. can oxidize a wide range of substrates (e.g., ethanol and butyrate) and transfer electrons to extracellular acceptors^47^. Their enrichment was particularly prominent under ethanol-supplemented conditions. However, in fluoroacetate-inhibited systems, although *Geobacter* remained abundant, acetoclastic methanogens such as *Methanothrix* were outcompeted by hydrogenotrophic methanogens, potentially due to disrupted TCA cycle activity of *Geobacter*^48^. Fluoroacetate likely impaired acetate oxidation in *Geobacter*, weakening its capacity to act as an electron-donating syntrophic partner.

This disruption by fluoroacetate may have led to the collapse of syntrophic acetate-linked electron flow. In the absence of inhibitors, only the ZVI particle biofilm condition sustained prolonged methane production without VFA accumulation, consistent with enhanced methanogenic activity observed in long-term reactors. In contrast, fluoroacetate treatment led to a sharp cessation of methane production and the accumulation of acetate and propionate, even in ZVI particle-inoculated reactors. Although fluoroacetate is commonly used to differentiate acetoclastic from hydrogenotrophic methanogenesis^17^, its impact in this context extended beyond classical acetoclastic methanogens. Transcript levels of *Methanothrix* were sharply suppressed, and other taxa potentially involved in acetate production or oxidation (e.g. SAO, SPO) were also affected. Despite the conductive ZVI surface, no improvement in methane accumulation was observed under fluoroacetate inhibition, implying that functional acetate-metabolizing partners were essential for the conductivity-dependent enhancement, whichever electron-transfer route was involved.

Predicted MetaCyc pathway analysis suggested that ZVI addition upregulated key pathways involved in VFA degradation, electron transport (Supplementary Results 3.5; Fig. S11). *Smithella* contributed to enhanced VFA degradation and was implicated in driving unfavored methanogenesis via DIET^42^. Meanwhile, *Geobacter* species were linked to upregulated pathways for benzoate and aromatic compound degradation (via benzoyl-CoA), nitrate reduction, and menaquinol biosynthesis, reflecting enhanced electron transport and respiratory flexibility in ZVI-enriched reactors^49^. These findings collectively indicate that short chain fatty acid (SCFA)-oxidizing syntrophs, rather than the classical SAO- or SPO-hydrogenotrophic partnerships alone, were central to the conductivity-dependent enhancement under psychrophilic conditions. Their enrichment on ZVI mirrors the trend reported for systems in which DIET has been demonstrated, but it remains a community-level signature.

Nevertheless, several limitations must be acknowledged. The evidence presented remains correlative: while community shifts and predicted pathway enrichment support syntrophic SCFA oxidation-mediated DIET, direct confirmation is lacking. This study relied on amplicon sequencing and function prediction; future research using metagenomics and metatranscriptomics will be essential to validate DIET-relevant gene expression and link them to specific SCFA oxidation.

## 5. Conclusion

This study shows that conductive ZVI particles enhance low-temperature anaerobic digestion of waste activated sludge by supporting a particle-associated syntrophic biofilm. The enhancement is conductivity-dependent: it was lost on non-conductive ZVI@SiO_2_ control, indicating that the dominant driver is a property of the native ZVI surface that is removed by electrical insulation, most plausibly its conductivity, rather than biofilm retention alone. The taxa enriched and the transcriptional responses on ZVI follow the trends reported for systems in which DIET has been demonstrated; DIET is therefore a plausible, but not directly demonstrated, contributor to this conductivity-dependent fraction. Compared to non-conductive ZVI@SiO_2_, ZVI supported the enrichment of syntrophic bacteria (*Smithella*, *Syntrophomonas*) and hydrogenotrophic methanogens, with elevated *mcrA* gene expression on particle surfaces. Functional analysis with methanogenic inhibition revealed upregulation of pathways linked to VFA degradation, electron transport metabolic capacity. These findings highlight the potential of conductive materials like ZVI to improve low-temperature AD performance and microbial resilience. From a practical standpoint, ZVI is an inexpensive, magnetically recoverable additive that was retained for 60 d with minimal wastage, and the 21–28% gain in methane production at 20 °C recovers part of the kinetic penalty that otherwise accompanies the energy savings of unheated digestion.

## Acknowledgements

We thank the Facility for Electron Microscopy Research at McGill University for technical support. We also gratefully acknowledge the operators and staff of the La Prairie wastewater treatment plant for granting site access and assisting with sample collection. This work was supported by a Natural Sciences and Engineering Research Council (NSERC) of Canada Collaborative Research and Development Grant (CRDPJ 500865-16) in collaboration with Air Liquide Canada.

## CRediT authorship contribution statement

**Chenxiao Liu**: Conceptualization, Methodology, Investigation, Data curation, Formal analysis, Visualization, Writing – original draft. **Dominic Frigon**: Conceptualization, Funding acquisition, Resources, Supervision, Project administration, Writing – review & editing.

