## Supplementary Information for "Boosting Methane Production at Low Temperatures: The Role of Zero Valent Iron-Driven Syntrophy in Anaerobic Digestion"

**Supplementary Methods**

*2.1. Synthesis of silicon dioxide coated zero valent iron particle*

*2.1.1 Synthesis protocol*

Iron powder (Fisher Scientific, Catalog No. AA0017014, spherical, <20 μm, 99.5%), acetone (Fisher Scientific, 99.9%), cyclohexane (Sigma-Aldrich, 99.8%), Triton^TM^ X-100 (Sigma-Aldrich), n-Butanol (Sigma-Aldrich, 99.8%), ammonia solution (Supelco, 25% w/w), tetraethyl orthosilicate (TEOS) (Sigma-Aldrich, 99.0%) were used as received.

Synthesis method adopted from Wu, Sun, Lu, Cheng and Gao [1] with modifications. A total of 50 g of iron powder was washed three times with acetone to remove surface impurities. A microemulsion was then prepared by combining 67.6 mL of cyclohexane, 15.5 mL of Triton^TM^ N X-100 as the surfactant, and 17.6 mL of n-butanol as the oil phase with 0.4 mL of ammonia solution as the water phase, followed by stirring for 1 hour at room temperature. The cleaned iron powders and 4.6 mL of TEOS were gradually introduced at 200 rpm stirring speed, and the mixture was stirred vigorously at 800 rpm for 10 hours, maintaining a pH of 9 by adding ammonia as needed. Afterward, the emulsion was demulsified using a solution of 25 vol% distilled water and 75 vol% acetone, and the lower phase was filtered and rinsed thoroughly with distilled water. Finally, the product was dried under anaerobic chamber overnight to yield SiO_2_ coated iron particles (ZVI@SiO_2_).

*2.1.2. Method of focused ion beam extreme high resolution scanning electron microscope (FIB-SEM) to exam the silicon dioxide coated ZVI*

The sample was attached to an SEM stub using carbon tape and coated with a 4 nm layer of platinum to improve electrical conductivity. The FIB-SEM dual beam system (FEI Helios Nanolab 660) has electron and ion beams intersecting at a 52° angle, allowing for both electron-beam-induced secondary electron imaging (SEM) and ion-beam-induced secondary electron imaging (IB). To extract a specimen in cross-sectional geometry, the sample stage was tilted approximately 52° so that the ion beam could strike the material perpendicularly.

For the FIB milling procedure, the target region was first coated with a 2 μm thick platinum layer to protect it from damage during milling. The target region was then milled using a Ga ion beam at a high accelerating voltage of 30 kV, starting with a maximum beam current of 65 nA. The ion beam current was subsequently reduced to 21 nA, with final milling performed at 9.3 nA. The milled regions were imaged using secondary electrons and backscattered electrons at accelerating voltages of 2 and 10 kV, with a beam current of 0.4 nA. Energy dispersive X-ray (EDX) analysis was conducted using an Octane Ultra 100 mm² SDD with the TEAM 3D EDS Analysis System, operating at an accelerating voltage of 10 kV and a beam current of 0.4 nA.

*2.1.3. DC resistivity of packed powder beds*

The apparent DC resistivity of ZVI and ZVI@SiO_2_ powders was measured in a two-probe configuration. 1 mL luer-slip syringes (inner diameter 4.69 mm; cross-sectional area A = 17.28 mm^2^ ) were trimmed and packed with each powder, leaving L = 8.94 mm of powder between two axially inserted stainless-steel probes. DC resistance was read with a benchtop precision resistance meter (B&K Precision 2831E, USA). Apparent DC resistivity was computed as ρ = RA/L.

*2.3. Batch incubation with microbial pathway inhibitors*

**Table S1**

Summary of experimental conditions in the various inhibition batch reactors

| Inoculum | Inhibitor Added | Inhibitor Concentration |
| --- | --- | --- |
| 20 ℃ ZVI@SiO2 added bulk sludge | Fluoroacetate | 1 mM |
|  | Chloroform | 100 µM |
|  | None (Control) |  |
| 20 ℃ ZVI added bulk sludge | Fluoroacetate | 1 mM |
|  | Chloroform | 100 µM |
|  | None (Control) |  |
| 20 ℃ Control bulk sludge | Fluoroacetate | 1 mM |
|  | Chloroform | 100 µM |
|  | None (Control) |  |
| 20 ℃ ZVI@SiO2 added particle | Fluoroacetate | 1 mM |
|  | Chloroform | 100 µM |
|  | None (Control) |  |
| 20 ℃ ZVI added particle | Fluoroacetate | 1 mM |
|  | Chloroform | 100 µM |
|  | None (Control) |  |

*2.3.1. Electron and carbon balance*

An electron-equivalent and carbon balance were closed for each inhibitor/control reactor using measured ethanol, VFA, CH_4_, CO_2_, H_2_ and volatile suspended solids (VSS), with all chemical species tracked in mg-COD/L (1 g-COD = 0.125 mol e⁻) and gases converted at 20 °C, 1 atm. The total input pool for each reactor was ethanol added at t = 0 (40 mM) plus initial biomass from the inoculum: 78–90 mg-COD per bulk reactor and 28–43 mg-COD per particle reactor. Liquid working volume and headspace volume were updated at every sampling event to reflect sample removal (8 mL for bulk reactors over 38 d; 12 mL for particle reactors over 62 d), cumulative gas production was summed, and cumulative VFA and ethanol mass removed in samples were added to the endpoint pool. The balances closed as Equation S1 for electron, Equation S2 for carbon:

$$Ethanol + Initial biomass = \mathrm{CH}_{4}+ \sum\mathrm{VFAs}+ H_{2}+End biomass + Unaccounted (Eq.S1)$$

$$Ethanol-C + Initial biomass-C = \mathrm{CH}_{4}-C+\mathrm{CO}_{2}-C+\sum\mathrm{VFAs}-C+ End biomass-C + Unaccounted-C (Eq.S2)$$

### *2.5.* 16S rRNA and mcrA amplicon sequencing

PCR was performed on DNA samples using Q5 polymerase, while RNA samples underwent PCR using the LunaScript® Multiplex One-Step RT-PCR Kit (New England Biolabs, Whitby, Canada). The 16S rRNA gene V4 region was amplified using primers F515 (5’-GTG YCA GCM GCC GCG GTA A-3’) and R806 (5’-GGA CTA CNV GGG TWT CTA AT-3’). The *mcrA* gene was amplified using primers mlas (5’-GGY GGT GTM GGN TTC ACH CAR TA-3’) and mcrA-rev (5’-CGT TCA TBG CGT AGT TVG GRT AGT-3’) [2]. After PCR amplification, the products were cleaned with the QIAquick PCR Purification Kit (Qiagen, Canada) and quantified using the Quant-iT™ PicoGreen™ dsDNA Assay Kit (Life Technologies Inc., Canada)[3].

*2.6. Digital PCR*

*2.6.1. Method of qPCR and digital PCR for 16S rRNA and mcrA gene*

For both DNA and RNA extractions, 16s rRNA and *mcrA* gene have been amplified with qPCR to determine the dilution fold for digital PCR. The sample Ct (cycle threshold) value from qPCR should higher than 25 before running the dPCR. This ensures that, when the sample is transferred to dPCR, the partitions will contain a manageable number of target molecules, reducing the risk of oversaturation. DNA samples were using Luna Universal qPCR Master Mix (NEB) to amplify and quantify with following program setting, and RNA samples were using Luna Universal One-Step RT-qPCR Kit (NEB) to add a reverse transcription step at beginning.

**Table S2**

qPCR conditions for 16s rRNA and *mcrA* gene

|  | 16s rRNA | | | *mcrA* | | |
| --- | --- | --- | --- | --- | --- | --- |
|  | Temperature | Time | Cycles | Temperature | Time | Cycle |
| Reverse Transcription (Only for RT-qPCR) | 55 °C | 10 m | 1 | 55 °C | 10 m | 1 |
| Initial Denaturation | 95 °C | 2 m | 1 | 95 °C | 2 m | 1 |
| Denaturation | 95 °C | 15 s | 40 | 95 °C | 15 s | 40 |
| Annealing | 55 °C | 30 s | 40 | 58 °C | 30 s | 40 |
| Extension | 72 °C | 1 m | 40 | 72 °C | 1 m 20 s | 40 |
| Melt Curve Analysis | 60–95 °C | 0.5 °C increment every 5 s | 1 | 60–95 °C | 0.5°C increment every 5 s | 1 |

The 16S rRNA gene was used as a housekeeping gene to normalize the absolute quantification of the *mcrA* gene obtained from digital PCR, following the specified digital PCR conditions. The restriction enzyme EcoRI (0.25 U µL−1) was added to the PCR solution to the long DNA and RNA, after template addition but prior to loading the nanoplate, the samples were incubated at room temperature for 10 min.

**Table S3**

Digital PCR conditions for 16s rRNA and *mcrA* gene

|  | 16s rRNA | | | *mcrA* | | |
| --- | --- | --- | --- | --- | --- | --- |
|  | Temperature | Time | Cycles | Temperature | Time | Cycle |
| Reverse Transcription (Only for RT-dPCR) | 50 °C | 40 m | 1 | 55 °C | 10 m | 1 |
| Initial Denaturation | 95 °C | 2 m | 1 | 95 °C | 2 m | 1 |
| Denaturation | 95 °C | 15 s | 40 | 95 °C | 15 s | 40 |
| Annealing | 50 °C | 15 s | 40 | 57 °C | 15 s | 40 |
| Extension | 72 °C | 30 s | 40 | 72 °C | 35 s | 40 |
| Final Extension | 35 °C | 5 min | 1 | 35 °C | 5 min | 1 |
| Imaging 1 | | | | | | |
| Denaturation | 95 °C | 15 s | 20 | 95 °C | 15 s | 20 |
| Annealing | 50 °C | 15 s | 20 | 57 °C | 15 s | 20 |
| Extension | 72 °C | 30 s | 20 | 72 °C | 35 s | 20 |
| Final Extension | 35 °C | 5 min | 1 | 35 °C | 5 min | 1 |
| Imaging 2 | | | | | | |

The dPCR mixtures for quantifying 16s rRNA gene total (archea and *mcrA* gene from cDNA will be prepared using QIAcuity EvaGreen PCR Kit (Qiagen) according to the manufacturer’s instructions. For taking pictures after cycling, the exposure duration was set at 200 ms with a gain of 3.

**Supplementary Results**

*3.1. Silicon dioxide coating*

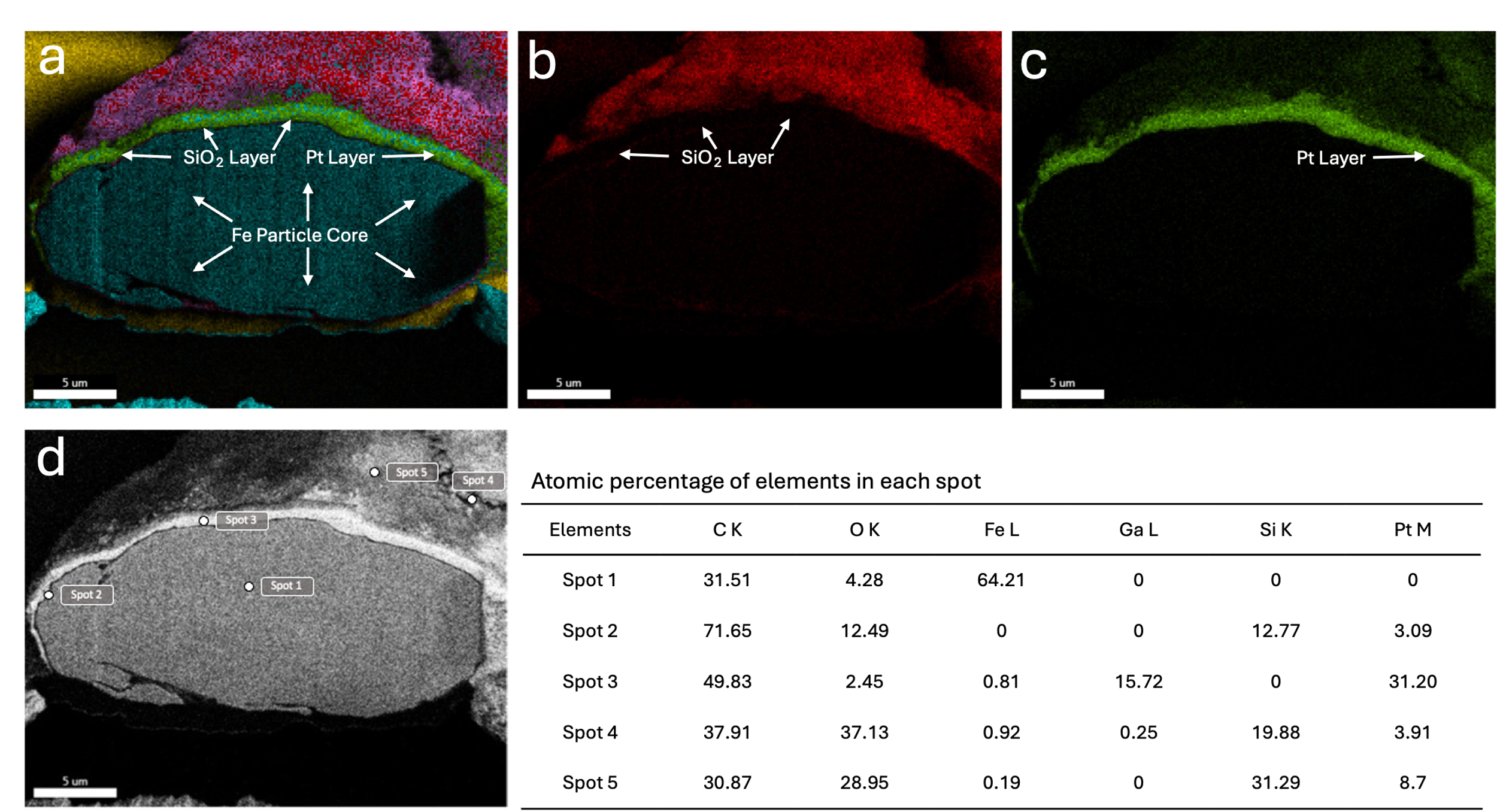

**Fig. S1.** EDS elemental mapping of ZVI@SiO₂ particles. a) EDS elemental mapping of all five selected elements: Fe (teal), Si (magenta), O (red), Pt (green), and C (gold). The carbon in the background represents the carbon tape used to attach samples for FIB-SEM. b) EDS elemental mapping of O (red); c) EDS elemental mapping of Pt (green); d) selected spots for elemental analysis and table of atomic percentage of elements in each spot.

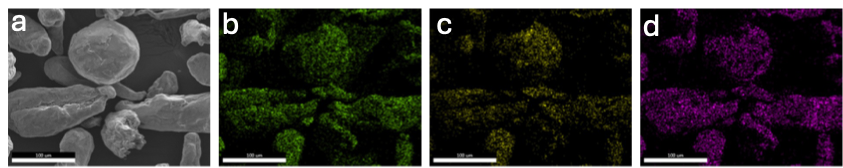

**Fig. S2.** SEM and EDS mapping of ZVI@SiO_2_ particles. a) SEM image of ZVI@SiO_2_ particles; b) EDS elemental mapping of O (green) with 62.88 atomic percentage; c) EDS elemental mapping of Fe (gold) with 10.88 atomic percentage; d) EDS elemental mapping of Si (magenta) with 26.23 atomic percentage.

**Table S4**

Apparent direct current resistivity of compacted ZVI and ZVI@SiO_2_ powder

| Material | R (Ω) | ρ (Ω·m) |
| --- | --- | --- |
| ZVI | 60 ± 10 | 0.116 ± 0.019 |
| ZVI@SiO_2_ | 2300 ± 128 | 4.44 ± 0.25 |

Note: R is electrical resistance, ρ is electrical resistivity.

*3.2. Long-term performances of digesters*

**Table S5**

Average performance of long-term anaerobic digesters at steady-state with ZVI@SiO_2_ and ZVI addition

| Reactor configuration |  | 20 ℃ | | |  | 35 ℃ | | | | |
| --- | --- | --- | --- | --- | --- | --- | --- | --- | --- | --- |
|  |  | Control | ZVI@SiO_2_ | ZVI |  | Control | | ZVI@SiO_2_ | | ZVI |
| pH |  | 7.1 ± 0.0 | 7.1 ± 0.1 | 7.0 ± 0.3 |  | 7.2 ± 0.0 | | 7.1 ± 0.1 | | 7.2 ± 0.1 |
| sCODout (mg/L) |  | 557 ± 31 | 572 ± 42 | 487 ± 12 |  | 483 ± 46 | | 456 ± 51 | | 433 ± 55 |
| Biogas production (mL/day) |  | 32 ± 1 | 32 ± 4 | 40 ± 1 |  | 44 ±3 | | 43 ± 4 | | 50 ± 3 |
| CH_4_% |  | 74 ± 2 | 76 ± 3 | 75 ± 1 |  | 68 ± 3 | | 68 ± 3 | | 68 ± 2 |
| CH_4_ production (mL/day) |  | 24 ± 1 | 24 ± 3 | 30 ± 2 |  | 30 ± 1 | | 29 ± 3 | | 34 ± 2 |
| SMPvss,in  (mL-CH_4_/g-VSS,in) |  | 143 ± 7 | 143 ± 15 | 183 ± 13 |  | 183 ±7 | | 177 ±12 | | 198 ± 15 |
| SMPvss,des  (mL-CH_4_/g-VSS,des) |  | 380 ± 26 | 382 ± 27 | 465 ± 25 |  | 451 ± 31 | | 456 ± 24 | | 480 ± 22 |
| VSSout (g/L) |  | 5.18 ± 0.06 | 5.17 ± 0.69 | 5.03 ± 0.22 |  | 4.20 ± 0.11 | | 4.39 ± 0.09 | | 4.46 ± 0.19 |
| VSS destruction (%) |  | 33 ± 3 | 33 ± 3 | 36 ± 2 |  | 45 ± 2 | | 46 ± 2 | | 47 ± 2 |
| VFAs (mgCOD/L) |  | 61 ± 2 | 58 ± 3 | 20 ± 1 |  | 15 ± 3 | 16 ± 4 | | 10 ± 2 | |
| Acetic acid (mgCOD/L) |  | 33 ± 2 | 26 ± 2 | 18 ± 1 |  | 13 ± 2 | 15 ± 4 | | 10 ± 2 | |
| Propionic acid (mgCOD/L) |  | 20 ± 1 | 16 ± 2 | 2 ±0 |  | 2 ± 2 | 1 ± 1 | |  | |
| Iso-butyric acid (mgCOD/L) |  |  | 8 ± 1 |  |  |  |  | |  | |
| Butyric acid (mgCOD/L) |  | 5 ± 1 | 6 ± 0 |  |  |  |  | |  | |
| Iso-valeric acid (mgCOD/L) |  | 3 ± 0 | 2 ± 0 |  |  |  |  | |  | |
| H_2_ production (mL/day) |  | 0 | 0 | 0 |  | 0 | | 0 | | 0 |

Note: Temperatures (20 ℃,and 35 ℃) indicate the digesters operational temperatures; Control has no particle addition, ZVI@SiO_2_ and ZVI represent the added particles in reactors; sCOD: Soluble chemical oxygen demand; SMPvss,in: Specific methane production using initial VSS; mL-CH_4_ : Volume of methane; g‑VSS,in : Mass of volatile suspended solid into the digester; SMPvss,des : Specific methane production using destroyed VSS; g-VSS,des : Mass of volatile suspended solid destroyed in the digester. ±: Standard deviation represents the variability across replicates and includes data from all measurements taken under steady-state conditions.

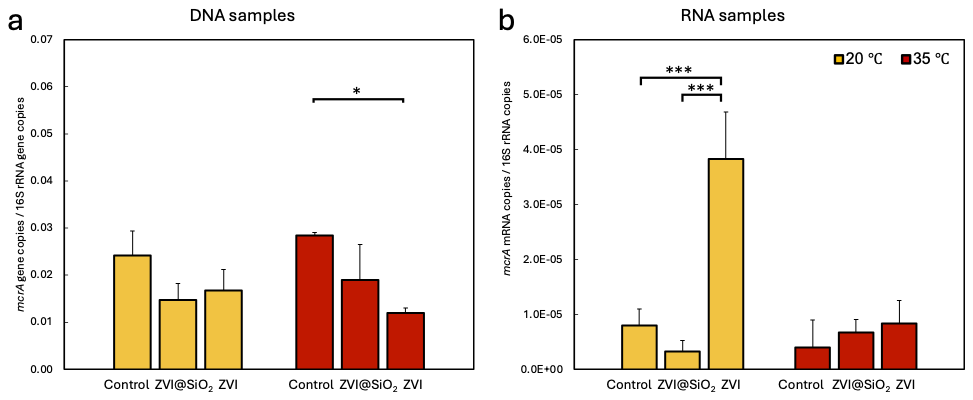

**Fig. S3.** Relative quantification of *mcrA* from bulk sludge normalized by their 16S rRNA at 20 °C and 35 °C. a) *mcrA* gene relative abundance in bulk from DNA samples; b) *mcrA* transcripts relative abundances in bulk from RNA samples. Significant differences (*, p < 0.05; ***, p < 0.001). Error bars represent standard deviations of biological replicates (n = 2 for non-particle controls, n = 4 for particle treatments).

*3.3. Microbial diversity of long-term reactors under different treatments*

*3.3.1. Alpha diversity of long-term anaerobic digesters*

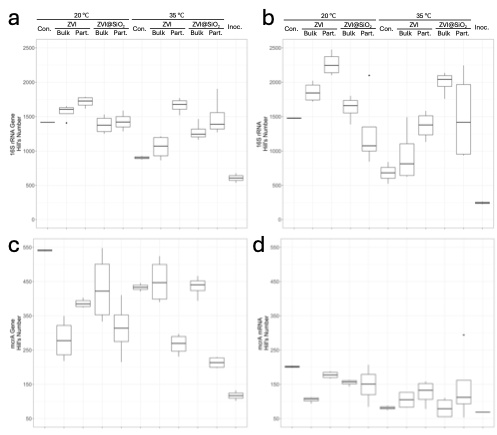

**Fig. S4.** Average first-order Hill’s number (exponential of Shannon index) for communities of anaerobic digestors receiving ZVI@SiO_2_ and ZVI particles at 20, and 35 °C a) 16S rRNA gene; b) 16S rRNA ; c) *mcrA* gene; d) *mcrA* mRNA. Con. was control reactor, Part. was particle, Inoc. was inoculum. Whiskers represent data within 1.5 times the interquartile range from the first and third quartiles; points beyond this range are plotted as outliers.

*3.3.2. Differences in bacterial population structures in long-term anaerobic digesters*

Sample beta-diversity and the most abundant bacteria genera are presented in Fig. S5a. For the total microbial community (Fig. S5a), temperature was the primary factor influencing sample clustering, with samples first grouping by temperature. Within each temperature group, sample location further differentiated communities, distinguishing microbial compositions on particle.

Temperature drove the total microbial community differences which was due to the favourable and adaptable of growth temperature. At 20 °C, the microbial community was characterized by the prevalence of *Christensenellaceae_R-7_group*, *Sedimentibacter* (both in phylum *Bacillota*, *DMER64*, *midas_g_12* (both in phylum *Bacteroidota*), *midas_g_249* (phylum *Synergistota*), and *midas_g_467* (phylum *Chloroflexota*). *Sedimentibacter* and *DMER64* were specifically enriched in bulk samples. At 20°C, particle-associated communities were enriched with Syntrophomonas, Cryptanaerobacter (both in phylum *Bacillota*), midas_g_762 (phylum *Synergistota*), and midas_g_3337 (phylum *Bacteroidota*). Within these, ZVI surface exhibited a further increase in Syntrophomonas and midas_g_762, along with the distinct enrichment of Syntrophorhabdus, and Smithella (both in phylum *Desulfobacterota*) which were specifically associated with ZVI treatment.

At **35 °C,** the microbial community showed elevated levels of midas_g_1237 (phylum *Acidobacteriota*) and midas_g_141 (phylum *Bacteroidota*). Exilispira (phylum Spirochaetota), SC103 (phylum Thermotogota), midas_g_141, and midas_g_33659 (phylum *Bacillota*) were specifically enriched in **bulk samples**. At 35 °C, **particle-associated communities** were enriched with Thermovirga (phylum *Synergistota*), midas_g_134 (phylum *Desulfobacterota*), midas_g_1469 (phylum *Chloroflexota*), midas_g_230 (phylum Caldatribacteriota), midas_g_326 (phylum *Bacteroidota*), and midas_g_5426 (phylum *Synergistota*).

Certain taxa were consistently **enriched in particle-associated communities** across both **20 °C and 35 °C**, highlighting their stable ecological role regardless of temperature. These included Smithella, Peptoclostridium (phylum *Bacillota*), Syntrophomonas, Syntrophorhabdus, and Syntrophus (phylum *Desulfobacterota*).

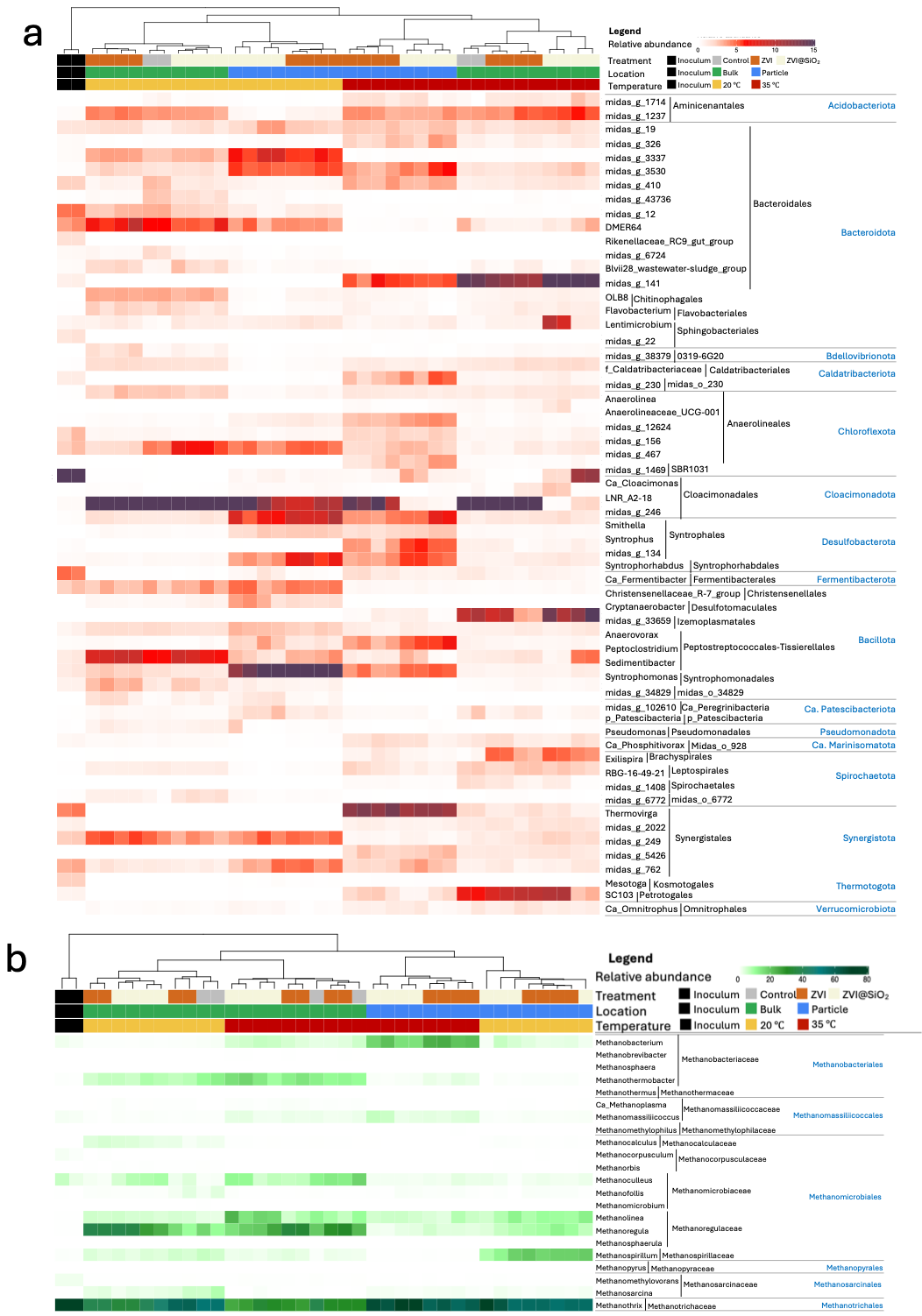

**Fig. S5.** Beta-diversity dendrogram and heatmap of the relative abundances of genera in anaerobic digestors receiving ZVI or ZVI@SiO_2_ at 20 or 35 °C determined by 16S rRNA gene (a) and mcrA gene (b) amplicon sequencing.

The dendrogram is based on Bray-Curtis dissimilarity calculated from amplicon sequence variants (ASVs) and clustered by UPGMA method. The heatmap includes the top 15 most abundant genera of each sample for both 16S rRNA gene and mcrA gene. For 16S rRNA gene, taxonomic annotations are shown from left to right as genus, order (after the vertical line), and phylum (in blue). For mcrA gene, annotations are shown as genus, family (after the vertical line), and order (in blue). All methanogenic genera shown for the mcrA gene panel belong to the phylum *Euryarchaeota*. Relative abundances are represented by color intensity, with red for the 16S rRNA gene and green for the mcrA gene. Reactor conditions are color-coded based on treatment (Control, Inoculum, ZVI, ZVI@SiO_2_), location (Bulk, Particle), and temperature (20 °C, 35 °C).

*3.3.3. Differences in active archaeal methanogenic population structures in long-term anaerobic digesters*

For the active methanogen community (Fig. S5b), samples first grouped by temperature. Within each temperature group, sample temperature further differentiated communities. Finally, treatment conditions played a key role in clustering.

At 35°C, *Methanomassiliicoccus* and *Methanobacterium* were more abundant overall, with *Methanobacterium* specifically enriched in particle-associated communities. In bulk sludge, *Methanoculleus* and *Methanothermobacter* were enriched. At 20°C, *Methanoregula* and *Methanolinea* were enriched in particle-associated communities, while *Methanoculleus* was more abundant in bulk sludge. Additionally, *Methanoregula* and *Methanolinea* were consistently enriched in bulk sludge at both 20°C and 35°C, indicating their presence across different thermal conditions. Regarding surface associations, *Methanothrix* was the most abundant methanogen on SiO_2_ surfaces at both temperatures, while *Methanospirillum* was more abundant on particles at 20°C but had low abundance in particle-associated communities at 35°C.

*3.4. Probing community food-web with methanogenic inhibitors*

*3.4.1. Performance of batch reactors*

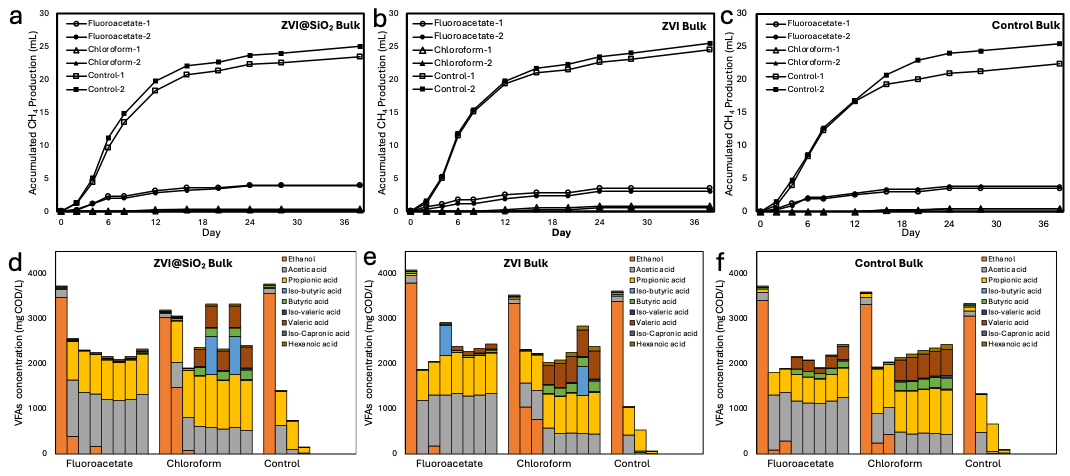

**Fig. S6.** Temporal accumulated CH_4_ produced (a, b, c) and VFAs concentration (d, e, f) profiles under various inhibitor additions in batch reactors at 20 °C inoculated with bulk from long-term digesters at the same temperature: (a, d) ZVI@SiO_2_ bulk, (b, e) ZVI bulk, (c, f) control bulk.

**Table S6**

Electron- and carbon-balance partition per reactor for the inhibitors and no-inhibitor treatments

| Inoculum | Inhibitor | CH_4_ | VFAs | H_2_ | Biomass | Closure% (e⁻) | CH_4_-C | CO_2_-C | VFAs-C | Closure% (C) |
| --- | --- | --- | --- | --- | --- | --- | --- | --- | --- | --- |
| ZVI@SiO_2_ Bulk | Fl | 4.5 | 38.5 | 0.2 | 38.9 | 82.1 | 2.8 | 0.5 | 45.1 | 96.5 |
|  | Ch | 0.3 | 40.3 | 0.2 | 36.9 | 77.8 | 0.2 | 0.8 | 44.3 | 90.9 |
|  | Con | 26.5 | 1.1 | 0.1 | 54.4 | 82.1 | 16.4 | 2.2 | 1.3 | 87.2 |
| ZVI@SiO_2_ Particle | Fl | 7.5 | 49.0 | 1.5 | 19.1 | 77.2 | 5.3 | 1.3 | 65.3 | 98.6 |
|  | Ch | 1.7 | 47.0 | 0.8 | 17.8 | 67.4 | 1.2 | 3.3 | 59.5 | 88.9 |
|  | Con | 11.4 | 43.1 | 0.3 | 19.6 | 74.3 | 8.0 | 1.5 | 59.3 | 96.2 |
| ZVI Bulk | Fl | 3.6 | 38.7 | 0.2 | 39.2 | 81.7 | 2.2 | 0.4 | 44.8 | 95.4 |
|  | Ch | 0.7 | 37.3 | 0.2 | 38.3 | 76.5 | 0.4 | 0.8 | 40.0 | 88.1 |
|  | Con | 25.8 | 0.8 | 0.1 | 55.1 | 81.7 | 15.8 | 2.2 | 0.9 | 86.4 |
| ZVI Particle | Fl | 6.6 | 51.3 | 0.9 | 21.1 | 79.9 | 4.5 | 1.7 | 66.9 | 102.0 |
|  | Ch | 0.9 | 37.5 | 0.3 | 25.6 | 64.3 | 0.6 | 1.8 | 45.2 | 82.6 |
|  | Con | 26.2 | 6.7 | 0.9 | 33.9 | 67.7 | 17.9 | 4.1 | 9.2 | 77.5 |
| Control Bulk | Fl | 4.1 | 38.8 | 0.2 | 41.1 | 84.1 | 2.5 | 0.5 | 44.3 | 97.9 |
|  | Ch | 0.5 | 38.9 | 0.2 | 36.3 | 75.8 | 0.3 | 0.7 | 41.9 | 87.5 |
|  | Con | 25.6 | 1.0 | 0.2 | 57.3 | 84.0 | 15.8 | 3.3 | 1.2 | 90.8 |

Inhibitor: Fl = fluoroacetate; Ch = chloroform; Con = no inhibitor. Values in percent of total-COD input (electron) or total-C input (carbon). Mean of 2 replicates.

The electron and carbon balance partition across batch experiment was given in Table S5. Bulk-reactor electron closures were 76–84 % across all treatments and carbon closures 86–98 %, indicating that ethanol-COD and initial biomass-COD were quantitatively partitioned between measured sinks with only modest residual attributable to dissolved bicarbonate and small septum losses over the 38-day incubation. Particle-reactor electron closures were lower (64–80 %) because the particle-reactor bulk liquid at endpoint was not sampled for suspended VSS.

*3.4.2. mcrA gene relative abundance and methanogenesis activity*

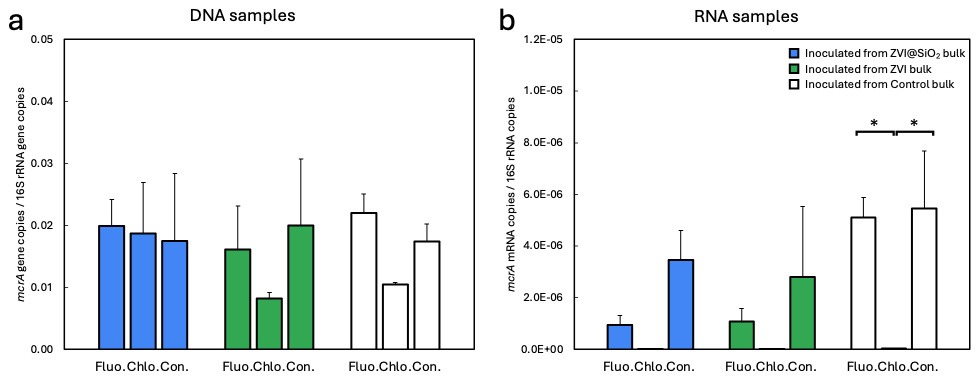

**Fig. S7.** Relative quantification of *mcrA* gene and mRNA from bulk sludge normalized by their 16S rRNA at 20 °C and 35 °C. a) *mcrA* gene relative abundance in bulk of inhibitors added reactors’ DNA samples; b) *mcrA* gene transcripts relative abundance in bulk of inhibitors added reactors’ RNA samples. Significant differences (*, p < 0.05). Error bars represent standard deviations of biological replicates (n = 2).

*3.4.3. Community diversity and structure of batch reactors*

Hill’s number analysis (Fig. S8) showed that chloroform-inhibited reactors had the lowest RNA-level alpha diversity across all inoculum types, while DNA-level diversity remained unaffected. Hill’s numbers for both 16S rRNA and mcrA mRNA (Fig. S8b, d) were consistently lower compared to their corresponding DNA extracts, indicating a smaller subset of transcriptionally active microbes, which may be consistent with the use of a single substrate (ethanol).

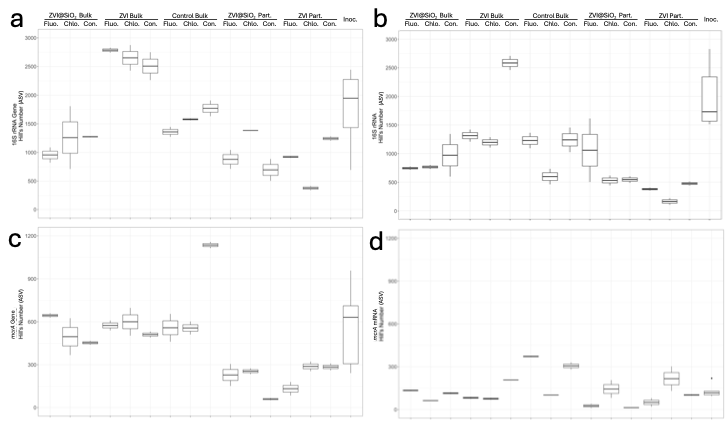

**Fig. S8.** Average first-order Hill’s number (exponential of Shannon index) for communities of anaerobic with inhibitors addition at 20 °C a) 16S rRNA gene; b) 16S rRNA; c) *mcrA* gene; d) *mcrA* mRNA.

Note: Con.= Control; Part. = Particle; Inoc. = Inoculum; Fluo. = Fluoroacetate; Chlo. = Chloroform

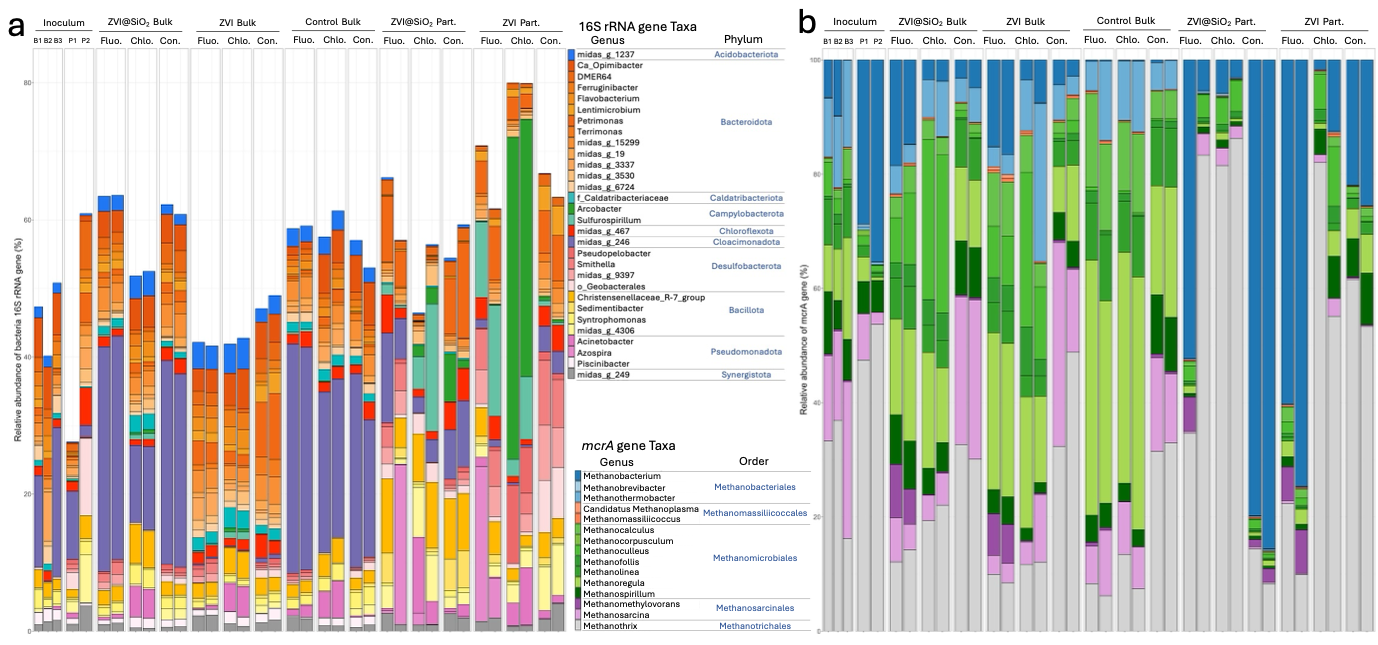

**Fig. S9.** Bar plot of the relative abundance of inhibitors added reactors at 20 °C (a) the 16S rRNA gene of top 30 bacteria genus and (b) the *mcrA* gene.

Inoculum was the initial inoculum from each reactor either bulk sludge or particle, B1 was the inoculum from ZVI@SiO2 bulk sludge, B2 was the inoculum from ZVI bulk sludge, B3 was the inoculum from control reactor bulk sludge, P1 was the inoculum from ZVI@SiO2 particle, P2 was the inoculum from ZVI particle. Con. was the control reactors without addition of inhibitor, Fluo. was fluoroacetate added reactors, Chlo. is chloroform added reactors. Taxonomic annotations are shown at the genus and phylum (in blue) levels for 16S rRNA gene, and at genus and phylum (in blue) levels for mcrA gene.

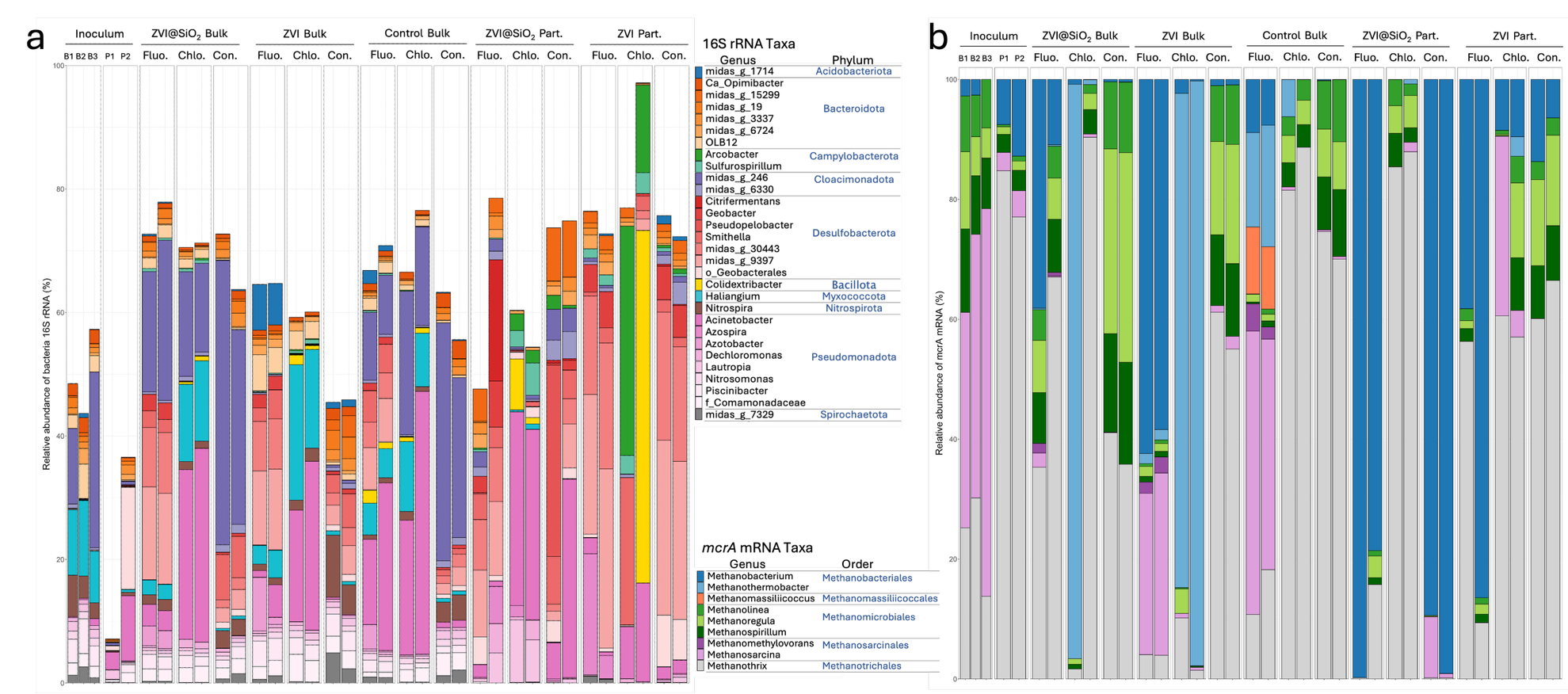

**Fig. S10.** Bar plot of the relative abundance of inhibitors added reactors at 20 °C (a) the 16S rRNA of top 30 bacteria genus and (b) the *mcrA* mRNA.

Inoculum was the initial inoculum from each reactor either bulk sludge or particle, B1 was the inoculum from ZVI@SiO2 bulk sludge, B2 was the inoculum from ZVI bulk sludge, B3 was the inoculum from control reactor bulk sludge, P1 was the inoculum from ZVI@SiO2 particle, P2 was the inoculum from ZVI particle. Con. was the control reactors without addition of inhibitor, Fluo. was fluoroacetate added reactors, Chlo. is chloroform added reactors. Taxonomic annotations are shown at the genus and phylum (in blue) levels for 16S rRNA, and at genus and phylum (in blue) levels for *mcrA* mRNA.

*3.5. Differential metabolic pathways*

The MetaCyc metabolic pathways were predicted from their 16S rRNA data of particle-associated communities to assess their potential abundances. Of these predicted pathways, the analysis suggested that 51 showed significantly higher potential abundances on ZVI particle-inoculated controls (no inhibitor added) than on ZVI@SiO_2_ particle-inoculated controls (Fig. S11a) and 31 were showed significant lower potential abundances on ZVI particle-inoculated controls (Fig. S11b). In ZVI particle-inoculated control reactors, 51 metabolic pathways were significantly upregulated, with nearly half (48%) associated with catabolic functions, including VFA degradation (14%), energy metabolism (20%), and aromatic compound breakdown (14%). Key potential pathways associated with VFA degradation included propanoate fermentation, pyruvate fermentation, phenylacetate degradation, and cinnamate and 3-hydroxycinnamate degradation. In terms of energy metabolism, six pathways for menaquinol biosynthesis were possibly more abundant, along with two nitrate reduction potential pathways. Several potential pathways related to aromatic compound degradation, including benzoyl-CoA degradation and toluene degradation, were also enriched. By contrast, in the 31 significantly downregulated pathways, the majority (68%) were involved in biosynthetic processes, with only small fractions linked to VFA degradation (3.2%) or energy metabolism (9.7%), underscoring that ZVI addition potentially enhanced catabolic metabolism.

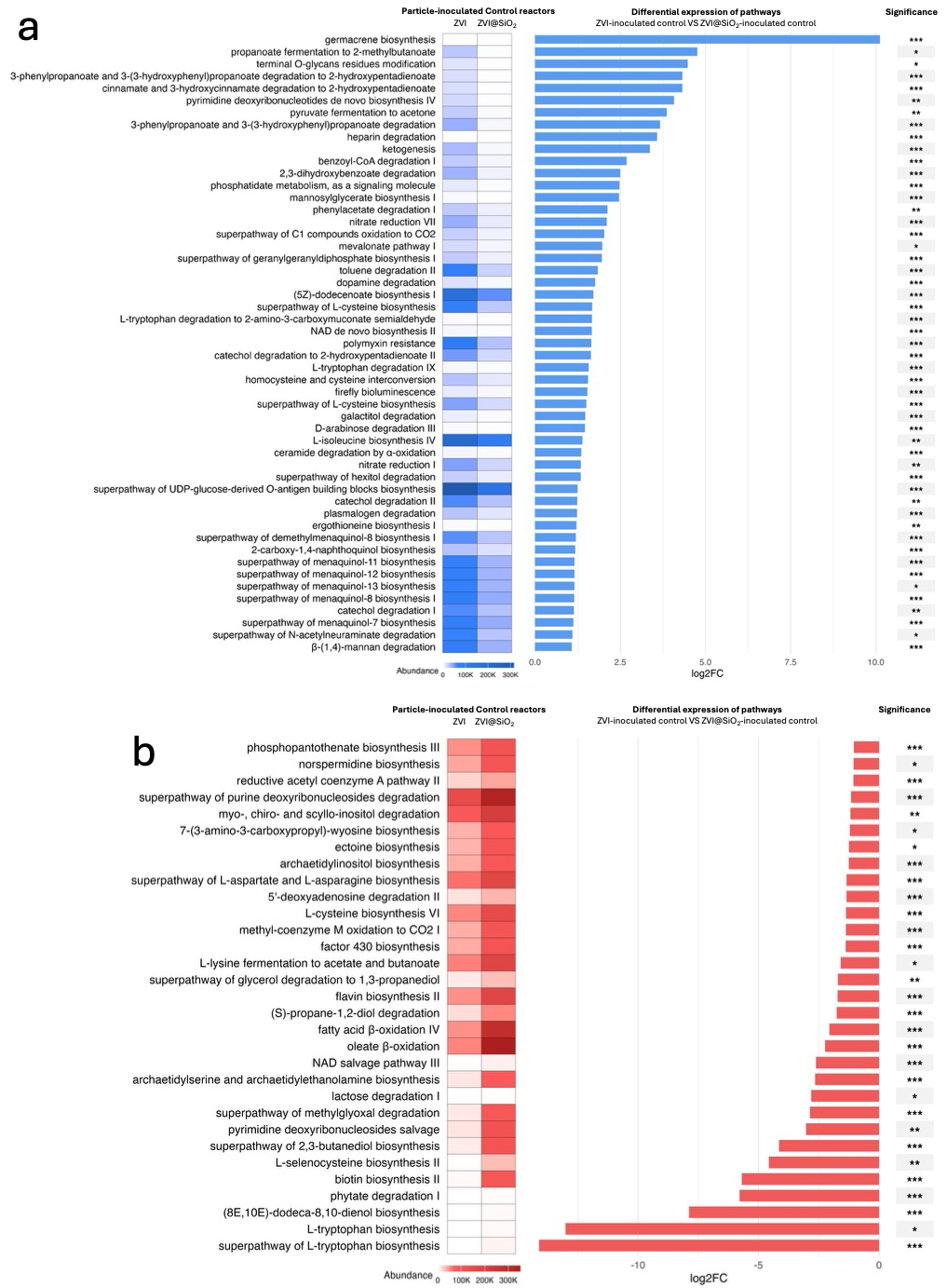

**Fig. S11.** MetaCyc metabolic pathways comparison between ZVI-inoculated control reactors and ZVI@SiO_2_-inoculated control reactors from RNA samples, a) significantly higher potential abundances; b) significantly lower potential abundances.

Heatmap is the abundance comparison. Bar plot shows the upregulated pathways for ZVI-inoculated control reactors, Wald test is used for the significance testing and adjusted P-values was used (*, p < 0.05; **, p < 0.01; ***, p < 0.001).

**Supplementary Discussion**

*4.3. Syntrophic interactions and functional adaptations in ZVI-amended anaerobic digestion*

*4.3.1. Active microbes selection from RNA to DNA ratio*

The biplots of RNA- and DNA-based genus abundances generally follow the 1:1 line (Fig. S12, S13, S14), while genera located above the 1:1 line indicate higher activity. At 20 °C, the particle surface biofilm revealed increased active members of *Smithella*, *Syntrophorhabdus*, and *Methanoregula* (Fig. S12a, b). When comparing each of their distance to the 1:1 line, indicating a larger ratio of RNA to DNA, the ZVI particle surface biofilm (Fig. S12a) had significantly further distances of these three genus then which on ZVI@SiO_2_ particle surface biofilm(Fig. S12b), emphasizing that ZVI further promoted their activity. Whereas they were below the 1:1 line in bulk and control communities (Fig. S14a-f). The improved activity comparing the RNA- and DNA-base methanogen abundances (*mcrA*) also showed further distance (0.1456) of *Methanoregula*, in addition of *Methanolinea*, above the 1:1 line at 20 °C ZVI particle surface biofilm (Fig. S12c) than it (0.0663) on ZVI@SiO_2_ particle surface (Fig. S12d), suggesting their distinctly active roles in the ZVI surface biofilm.

At 35 °C, the active microbes were *midas_g_1714* (order *Aminicenantales*), *Smithella*, *Syntrophorhabdus*, *midas_g_134* (order Syntrophales), and *midas_g_19* (family *Bacteroidetes vadinHA17*) on the particle surface biofilm (Fig. S13a, b). Excepting *midas_g_1714*, all the other active bacteria tended to have longer distance to the 1:1 line on ZVI@SiO_2_ particle surface. Comparing among *mcrA* RNA and DNA (Fig. S13c, d), *Methanoregula*, *Methanolinea*, along with *Methanoculleus* implied their activity greater on ZVI particle surface.

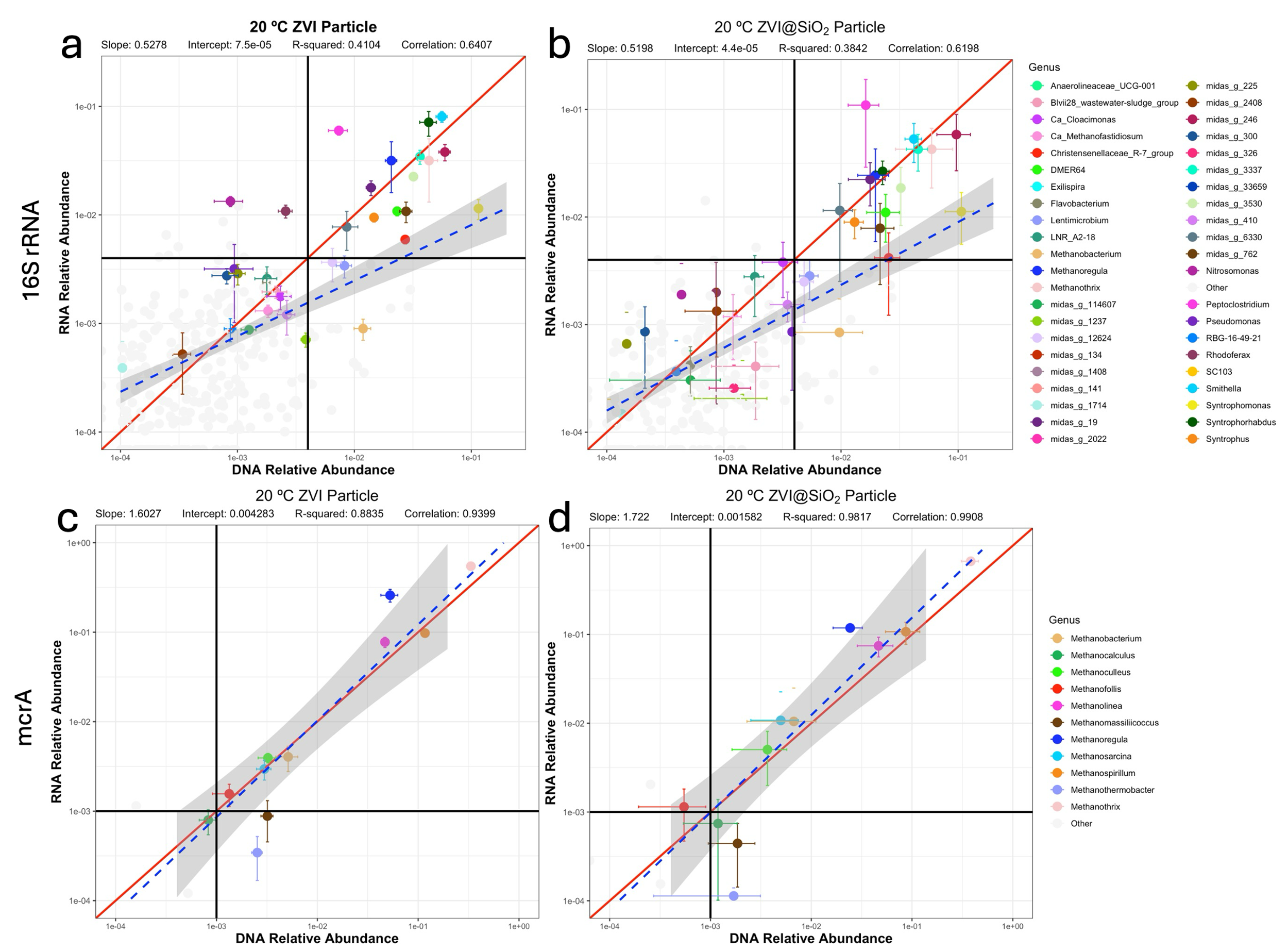

**Fig. S12.** DNA to RNA relative abundance biplot of 20 °C reactor particle biofilm at genus level a) ZVI particle biomass of 16S rRNA; b) ZVI@SiO2 particle biomass of 16S rRNA ; c) ZVI particle biomass of mcrA; d) ZVI@SiO2 particle biomass of mcrA. Red line is RNA equals to DNA, blue dash line is linear regression, crossed black solid lines are threshold for both DNA and RNA relative abundance at 10-3. Relative abundance is in log scale, error bars are calculated from both DNA and RNA sample replicates.

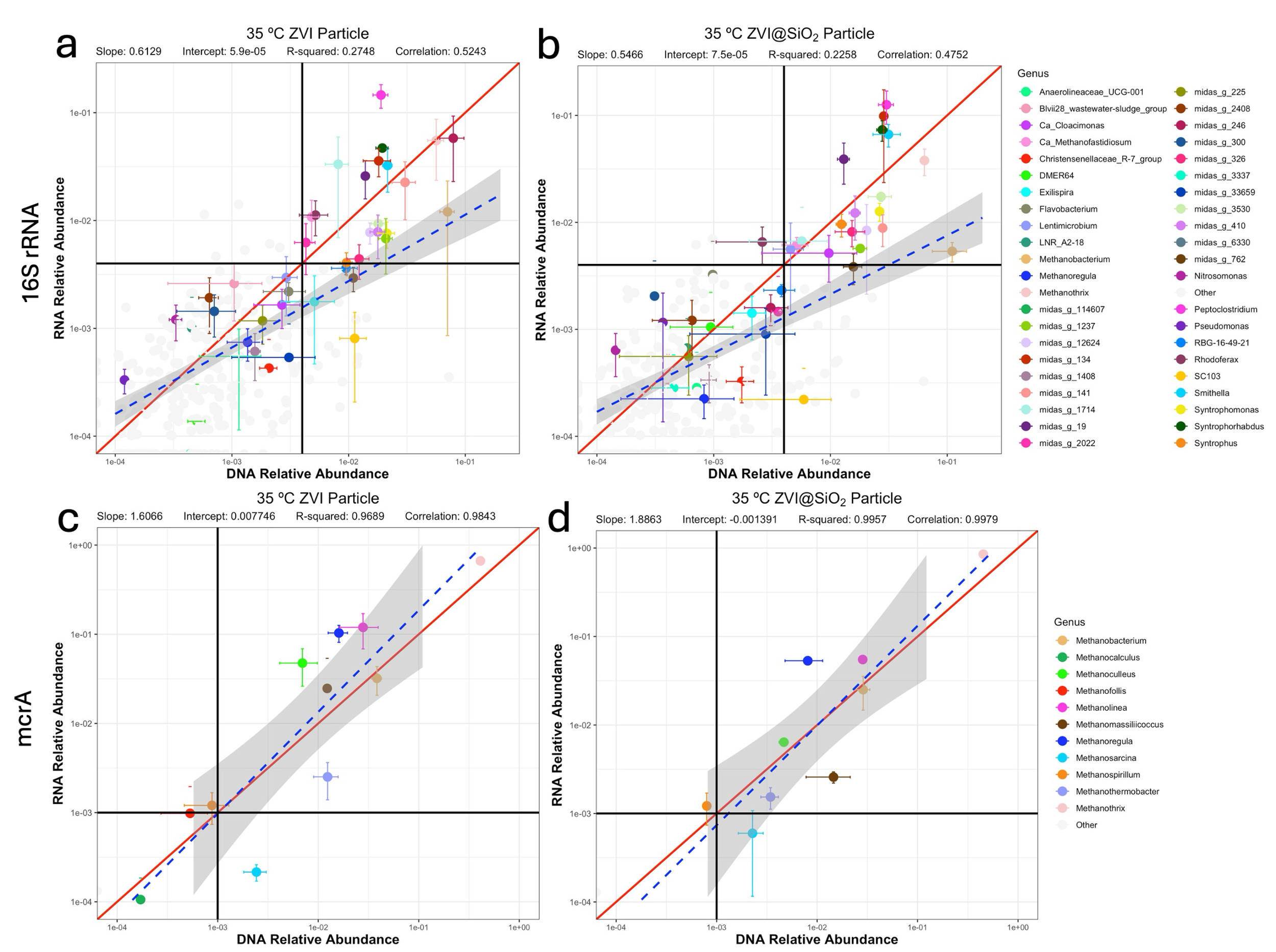

**Fig. S13.** DNA to RNA relative abundance biplot of 35 °C reactor particle biofilm at genus level a) ZVI particle biomass of 16S rRNA; b) ZVI@SiO_2_ particle biomass of 16S rRNA ; c) ZVI particle biomass of *mcrA*; d) ZVI@SiO_2_ particle biomass of *mcrA*.

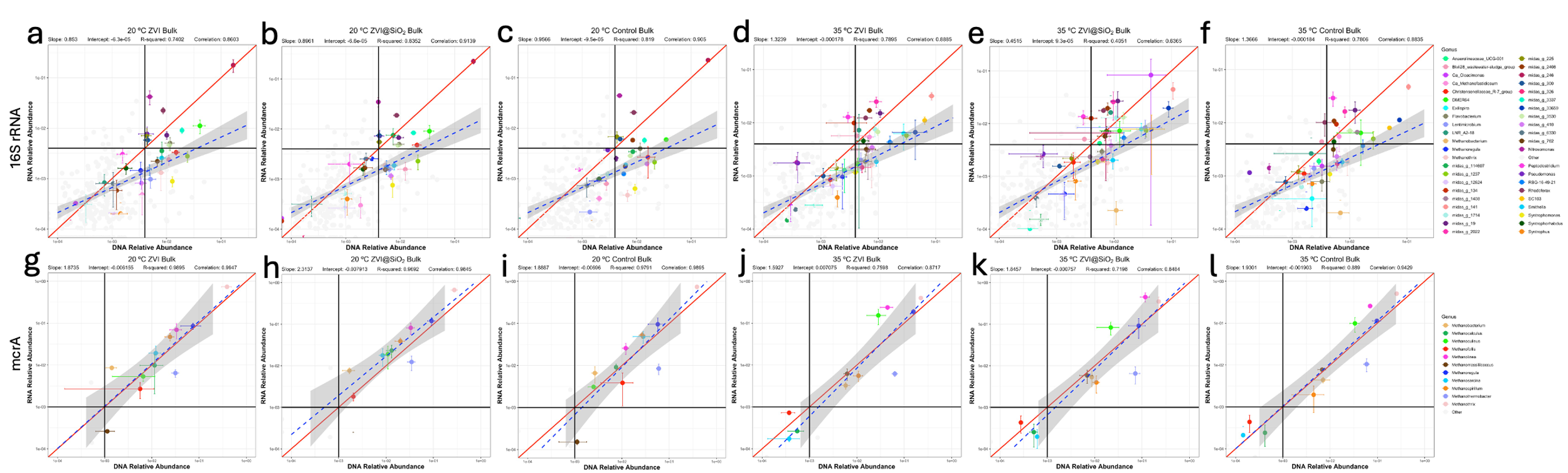

**Fig. S14.** DNA to RNA relative abundance biplot of 20 and 35 °C reactor bulk sludge at genus level. a) ZVI reactor bulk sludge of 16S rRNA at 20 °C; b) ZVI@SiO_2_ reactor bulk sludge of 16S rRNA at 20 °C; c) Control reactor bulk sludge of 16S rRNA at 20 °C; d) ZVI reactor bulk sludge of 16S rRNA at 35 °C; e) ZVI@SiO_2_ reactor bulk sludge of 16S rRNA at 35 °C; f) Control reactor bulk sludge of 16S rRNA at 35 °C; g) ZVI reactor bulk sludge of *mcrA* at 20 °C; h) ZVI@SiO_2_ reactor bulk sludge of *mcrA* at 20 °C C; i) Control reactor bulk sludge of *mcrA* at 20 °C; j) ZVI reactor bulk sludge of *mcrA* at 35 °C; k) ZVI@SiO_2_ reactor bulk sludge of *mcrA* at 35 °C C; l) Control reactor bulk sludge of *mcrA* at 35 °C.

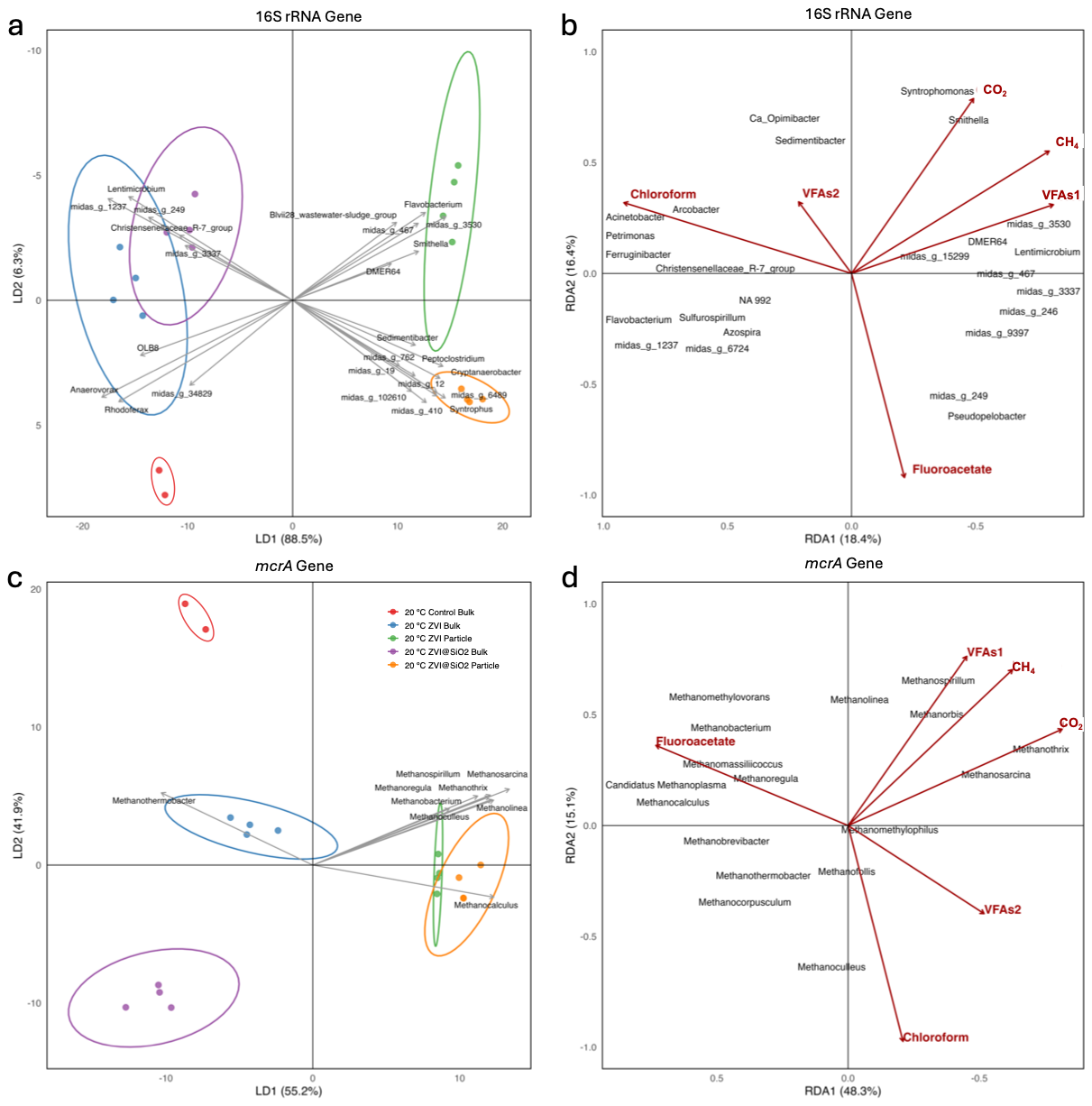

**Fig. S15**. Key microbial taxa at 20 °C determined by DNA extracts and differentiated by long-term reactor conditions (Canonical Discriminant Analysis (CDA) of 16S rRNA gene bacterial population [a] and *mcr*A gene for methanogens [c]) or short-term batch reactors performances with methanogenesis inhibitor additions (Redundancy Analysis [RDA] of 16S rRNA gene [b] and *mcr*A gene [d]).

For the RDA, the top 30 genera identified in digesters were included in the analyses. The red arrows represent different performance data and inhibitor additions binary coded as 0/1; note that the arrow no addition control reactors would point along the CO_2_ arrows. Temporal profiles of concentrations in VFAs were reduced in dimensions into two factors together explaining 26% of variability by principal component analysis using Euclidean dissimilarities.

1. Note on Section Numbering:

   This document follows the same section numbering as the main text to ensure clear and consistent cross-referencing. All supplementary information is organized according to the corresponding sections in the main text.

   For example, supplementary method related to Section 2.1. Synthesis of silicon dioxide coated zero valent iron particle in the main text are addressed under the same heading "2.1. Synthesis of silicon dioxide coated zero valent iron particle " in this document.
